# Experimental Design Optimization for Brain Microstructure Imaging via Automatic Differentiation

**DOI:** 10.1101/2024.12.09.627646

**Authors:** Hamza Farooq, Yongxin Chen, Ghulam Rasool, Tryphon T. Georgiou, Christophe Lenglet

## Abstract

Accurate characterization of the brain microstructure using diffusion MRI (dMRI) relies on optimal (i.e. information-rich) scanning protocols. Presently, without such protocols, extensive datasets are necessary to image the intricate microarchitecture of the brain by fitting complex biophysical models to the data. This impedes the full realization of dMRI’s potential, particularly in time-constrained clinical scans. To tackle this, we introduce a novel framework based on the Cramér-Rao lower bound (CRLB) to optimize dMRI protocols for any multi-compartment biophysical models, to accurately capture features like axonal diameter index and multiple fiber orientations in a voxel. Unlike previous methods, limited in model complexity or parameter scope, our framework handles all model parameters, including water diffusivities, enhancing estimation fidelity. Leveraging automatic differentiation and parallel computing via TensorFlow, this approach systematically explores the entire parameter space for comprehensive scanning protocol optimization. By optimizing dMRI protocols, this work stands to significantly enhance biophysical modeling accuracy, thereby deepening our understanding of the brain microstructure in health and disease.

## Introduction

Diffusion magnetic resonance imaging (dMRI) is a non-invasive neuroimaging technique that probes the microstructural properties of biological tissues by quantifying the Brownian motion of water molecules. This technique offers significant potential for investigating subtle alterations in tissue architecture associated with various neurological and pathological conditions (1).

To extract quantitative information about these microstructural features, dMRI data undergoes analysis to estimate biophysical models. These models are mathematical frameworks that incorporate the influence of cellular components, such as axons and myelin sheaths, on the diffusion process. By fitting biophysical models to dMRI data, researchers can estimate a range of tissue properties, providing valuable insights into tissue health and function.

Biophysical models depict tissue microstructure in greater and more specific details than simpler models by incorporating separate compartments for intra-axonal, extra-axonal, and cerebrospinal fluid (CSF). However, these complex models heavily rely on the specific parameters chosen during the dMRI acquisitions (e.g., pulse sequence, diffusion time, gradient strength, etc.) for accuracy. The challenge therefore lies in selecting optimal scanning parameters to ensure the most accurate estimation of these model parameters. In addition, accurate estimation of the parameters of the biophysical model requires the acquisition of multiple dMRI images with varying scanner parameters, which can significantly extend the scan time. In vivo studies and clinical applications face strict limits on total scan time due to factors like participant comfort and artifacts. Therefore, it is crucial to optimize the dMRI acquisition protocols to ensure that the most informative data is collected within a practical timeframe.

This work focuses on the optimization of dMRI acquisition protocols for the estimation of complex, multi-compartment biophysical models in the context of in vivo studies with limited scan times. The fundamental question we address is: How can one design dMRI experiments to acquire the most informative signal within clinically feasible time constraints, while employing a framework that leverages the information from multiple sophisticated biophysical models for data analysis?

Minimizing the Cramér–Rao lower bound (CRLB) of the tissue biophysical model parameters can be used to discover optimal dMRI data acquisition schemes. The method identifies a set of protocol parameters that minimizes the partial derivatives of the model parameters, with the advantages that it considers all possible combinations of scanning protocols and that it can be applied during the experiment planning before acquiring any data. Previous work has used the CRLB to optimize diffusion acquisition schemes for relatively simple biophysical models e.g., a simplified CHARMED model (2; 3), optimization of diffusion measurements (4), bi-tensor model (5), Diffusion Kurtosis Imaging (DKI) (6), optimizing quantitative sequences for Bloch simulations (7), Stick-and-Zeppelin model (8), MR finger printing (9), and q-space trajectory imaging using diffusion tensor distribution (10).

In contrast to the CRLB framework, data-driven approaches offer an alternative method for identifying optimal scanning protocols for multi-compartment biophysical models. For instance, a recent study (11) demonstrates this approach by acquiring q-space shells at various b-values and with very high angular resolutions, and iteratively evaluating subsets of these shells. Results from these subsets were compared with the full dataset, aiming to pinpoint the most effective protocol achievable within scanning time constraints. However, it is important to note that even the full dataset might still yield inaccurate model parameter estimates due to fitting errors. These inaccuracies could potentially skew the determination of the optimal protocol. On the other hand, CRLB-based methods are immune to such estimation errors, providing a more reliable approach for protocol optimization.

While previous studies have addressed the optimization of MR protocols (with or without CRLB), their scope was often limited in three key aspects: 1) Prior studies frequently employed simplified biophysical models, primarily due to difficulty in calculating derivatives of more intricate multi-compartment models; 2) The protocol optimization efforts were often restricted to specific model parameters, such as axonal fiber orientations, while fixing other parameters like water diffusivity values which may affect the overall optimization process; 3) The nonconvex nature of the optimization problem often necessitated the use of heuristic optimization algorithms. While these algorithms can identify reasonable solutions, they cannot guarantee finding the globally optimal acquisition protocol. This limitation could lead to suboptimal data collection strategies, potentially hindering the accuracy of tissue models parameter estimation.

The proposed solution is a generalized framework that can handle complex, multi-compartment tissue models. This framework allows for optimization across all model parameters, encompassing those related to tissue geometry including water diffusivity. Specifically, we aim to overcome the limitations of previous efforts to find optimal scanning protocols for multicompartment biophysical models by: 1) Extending our focus to any multi-compartment biophysical models capable of detecting intricate brain tissue features like axon diameter, distribution, and multiple fiber orientations; 2) Optimizing over all model parameters relating to tissue geometry, including water diffusivity values, to enhanced model parameter estimation fidelity; 3) Leveraging differentiable programming in TensorFlow (12), we employ derivative-based Fisher information matrix (FIM) calculation methods, enabling efficient optimization through automatic differentiation and parallel computing with GPUs; 4) The proposed approach systematically explores the entire parameter space of the acquisition protocol, and also span extreme ranges of biologically feasible tissue geometrical features, to ensure comprehensive optimization. Fig. 1 summarizes the overall workflow of the proposed framework.

**Figure 1.**
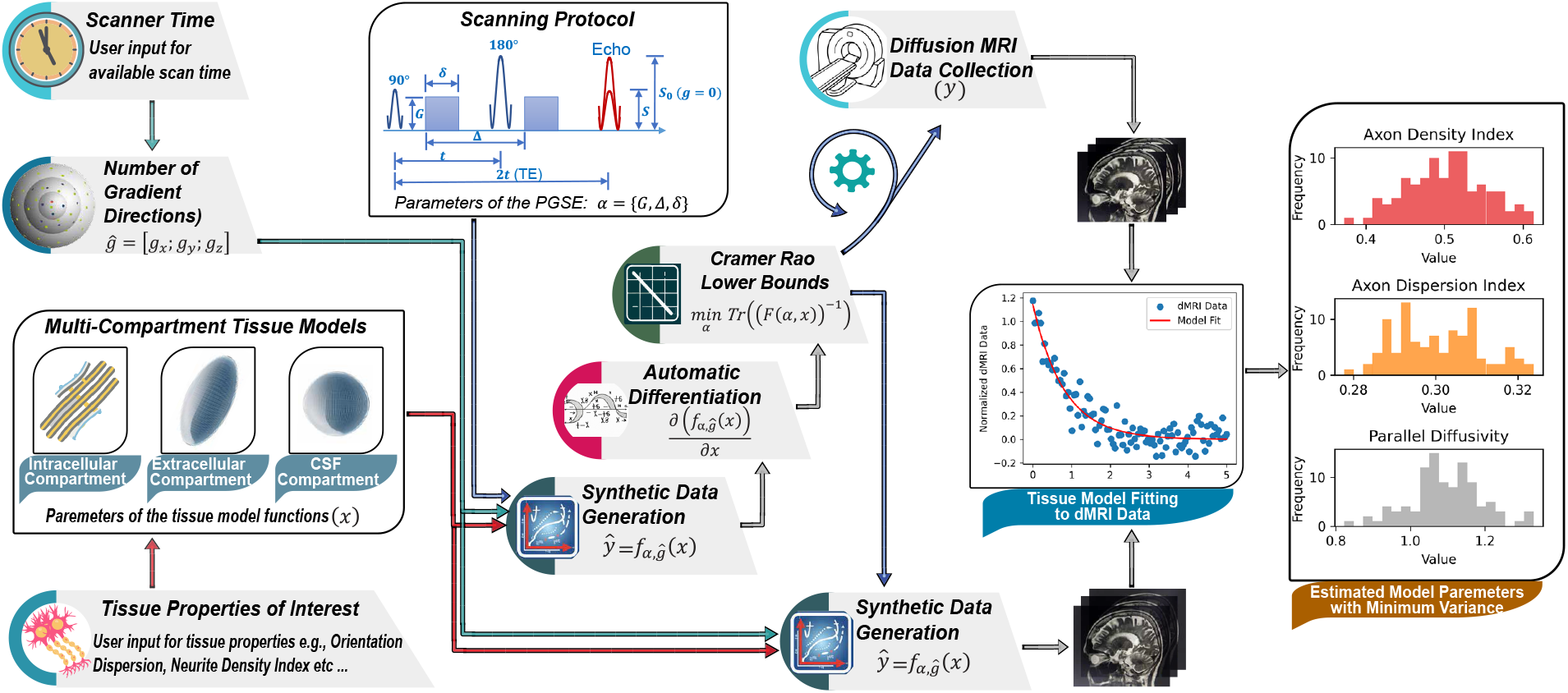
The figure illustrates a framework for optimizing diffusion MRI (dMRI) scans using Pulse Gradient Spin Echo (PGSE) sequences. The objective is to acquire the most informative dMRI signal for specific biophysical models, within a constrained scan time. This optimization process involves: 1) User-defined selection of tissue properties of interest (biophysical model selection) and available scanning time, 2) Generation of synthetic dMRI data using the model, followed by computation of the Cramér-Rao lower bound (CRLB) with automatic differentiation to determine the optimal scanning protocol parameters (*α*). 3) The process involves iterative testing of the candidate scanning parameters (considering scanner hardware limitations and balancing information gained from the signal and SNR), and finally, 4) Fitting of the model to dMRI data, resulting in estimated model parameters (*x*) with minimal variance for reliable microstructure imaging.

By providing a method to optimize dMRI acquisition protocols for complex models, this research has the potential to significantly improve the accuracy of biophysical modeling in dMRI. This, in turn, could lead to a better understanding of the properties of brain tissue in both healthy and diseased states. The subsequent sections of the paper are structured as follows: The Introduction is followed by the Results section, which details the optimal scanning parameters identified through our proposed framework. The subsequent Methods section then focuses on the framework itself, explaining its details, the model parameters it considers, and the scanning protocol parameters it optimizes. Following this, the Discussion section explores the broader context of these findings by relating them to existing research. Finally, the paper concludes with a concise summary of the key takeaways from this work.

## Results

This section aims to first establish the reliable recovery of water diffusivity values in brain tissue by optimizing dMRI scanning parameters. We achieve this by optimizing the dMRI scanning parameters in four distinct biophysical models: Diffusion Tensor (13), Neurite Orientation Dispersion and Density Imaging (NODDI) (14), ActiveAx (15), and Ball-and-Three-Sticks (16). Subsequently, we analyze each model, encompassing all model parameters, including diffusivity values, to identify the optimal overall combination of scanning parameters. Also, we revisit optimal multi-shell protocols for NODDI (14) and ActiveAx (15) models from previous studies, using the same parameter constraints. This enables us to confirm that our method finds consistent results with previous studies when placed under similar constraints.

### Understanding the impact of scanning parameters on water diffusivity estimation

Fig. 2 illustrates the relationship between water diffusivity and the optimal b-values (refer to Methods section for details on b-values/ scanning parameters) which is essential for the estimation of diffusivity values with minimal variance (i.e., minimum Cramér-Rao lower bound, CRLB). Each panel (A-D) corresponds to a different biophysical model. All sub-figures plot the magnitude of the gradient value G on the x-axis (*mT*.*m*^−1^) and the log-scale CRLB on the y-axis. The color scheme signifies different diffusivity components: red represents the primary/intrinsic diffusivity along the main fiber/axonal orientation (*λ*_1_ or *λ*_‖_), blue depicts the diffusivity along the secondary main orientation (*λ*_2_ or *λ*_⊥_), and green corresponds to the tertiary diffusion direction (*λ*_3_). Details for each model is as follows:

**Figure 2.**
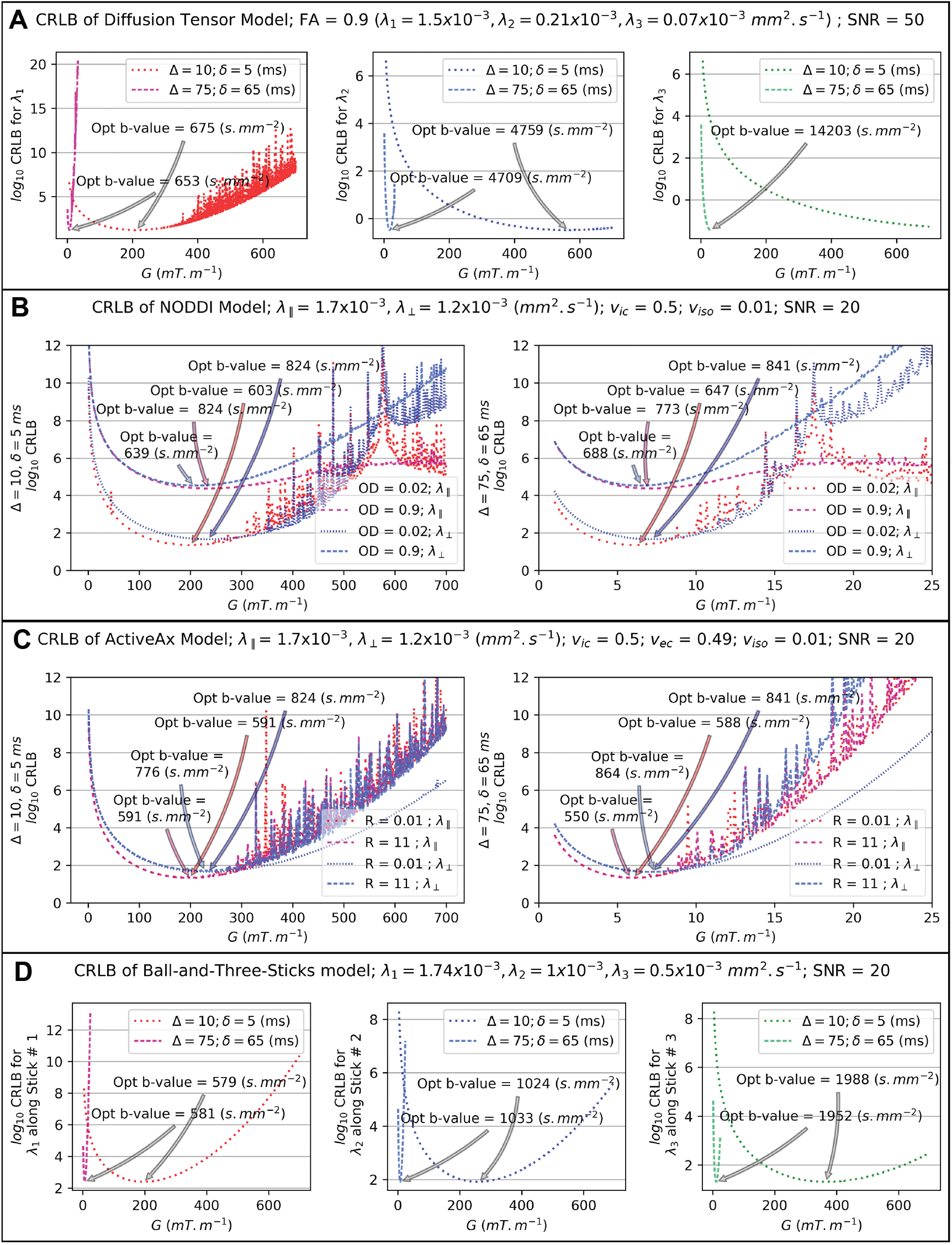
The figure shows the optimal b-values required for estimation of diffusivity values with minimal variance (i.e., minimum CRLB for diffusivities). Each panel (A-D) corresponds to a distinct tissue biophysical model: Diffusion Tensor (13), NODDI (14), ActiveAx (15), and Ball-and-Three-Sticks (16). respectively. All sub-figures use the magnitude of the MR system gradient values G on the x-axis (mT.m-1) and the CRLB on the y-axis using a logarithmic scale. The color scheme signifies different diffusivity components: red represents the primary/intrinsic diffusivity along the main fiber/axonal orientation (*λ*_1_ or *λ*_‖_), blue depicts the diffusivity along the secondary main orientation (*λ*_2_ or *λ*_⊥_), and green corresponds to the tertiary diffusion direction (*λ*_3_).

#### Diffusion Tensor

Fig.2A addresses the diffusion tensor model (13), a simple yet widely used approach for dMRI-based microstructure evaluation. The model incorporates six parameters, including three mutually perpendicular eigenvectors that represent the primary diffusion directions within a voxel (see Supplementary Fig. S1 for details). Human brain diffusivity values typically range from 0.02 to 1.74 x 10^−3^ *mm*^2^.*s*^−1^ (see (17) and references therein). To reflect this range, we employed a diffusion tensor with Fractional Anisotropy (FA) = 0.9, with three distinct eigenvalues within the human brain’s feasible range. For clarity, we used only three diffusion gradient directions perfectly aligned with the model’s eigenvectors and an ideal SNR of 50.

The three sub-figures in Fig. 2A depict the CRLB variations for each eigenvalue, revealing an optimal b-value around the reciprocal of the diffusivity value. The finding supports studies comparing Apparent Diffusion Constant (ADC) with b-values. For instance, Mukherjee et al. (18; 19) suggested an optimal b-value defined by the inverse of the tissue ADC.

These findings underscore the dependence of optimal scanning protocols on specific diffusivity values. Estimating multiple brain tissue diffusivities necessitates protocols incorporating multiple b-values. The challenge of estimating lower diffusivities is evident, as they require high b-values for optimal CRLB. For example, in Fig.2A (right sub-figure), a diffusivity of 0.07 x 10^−3^ *mm*^2^.*s*^−1^ necessitates a b-value of approximately 14 x 10^3^ *s*.*mm*^−2^ to minimize the CRLB. Specifically, achieving the CRLB in the diffusion tensor model depends solely on the b-value, independent of specific *G* (*mT*.*m*^−1^), Δ (ms), and *δ* (ms) combinations (refer to Methods section for scanning parameters description and Fig. S1). Additionally, these results maintain their consistency irrespective of the selections made for the rotation parameters of the diffusion tensor, namely, *θ, ϕ*, and *β* (rad).

#### NODDI

The model offers valuable insights into white matter integrity and pathology. It leverages a three-compartment tissue model to separately analyze intricate intra- and extracellular properties. NODDI can characterize two key aspects of axonal pathology: axonal packing density within white matter, quantified by the intracellular volume fraction (*v*_*ic*_), and the spatial dispersion of axons, captured by the orientation dispersion index (*OD*). Prior research has shown the influence of diffusivity variations on NODDI parameter estimates (20; 21).

The results presented here remain consistent regardless of the specific diffusivity values chosen, though we adopted a range from 0.5 to 1.74 x 10^−3^ *mm*^2^.*s*^−1^ to align with prior literature (14; 20). The analysis employed two perfectly aligned gradient directions, one along the fiber orientation and the other perpendicular, corresponding to parallel and perpendicular diffusivities. Additionally, we examined the impact of *OD* on diffusivity estimation using extreme values of 0.2 and 0.9. To ensure a balanced contribution of the signal from the intra- and extracellular compartments, the intracellular volume fraction (*v*_*ic*_) was fixed at 0.5.

In Fig.2, panel B shows the minimum achievable CRLB for various intrinsic/parallel and perpendicular diffusivity values. The two sub-figures in Panel B correspond to short (Δ = 10 ms, *δ* = 5 ms) and long (Δ = 75 ms, *δ* = 65 ms) diffusion time/pulse duration parameters of the Pulse Gradient Spin Echo, PGSE (please see Methods section for the details). The results reveal that optimal b-values, marked by minimal CRLB, are roughly reciprocal to diffusivity values, irrespective of Δ and *δ* choices. However, *OD* values affect optimal CRLB or b-values. In regions with minimal axonal dispersion (e.g., *OD* = 0.02 meaning high anisotropy), both intrinsic and perpendicular diffusivities achieve optimal CRLB at b-values roughly reciprocal to their respective diffusivity values. Conversely, under high orientation dispersion (e.g., *OD* = 0.9), indicative of more complex diffusion patterns, parallel diffusivities necessitate relatively higher b-values, while perpendicular diffusivities achieve optimal CRLB at relatively lower b-values. This observation aligns with the anticipated lower parallel diffusivity and higher perpendicular diffusivity in complex (e.g. fiber crossings) diffusion scenario. Similar results, concerning lower diffusivity values such as those found in gray matter and the infant brain, are depicted in Supplementary Fig. S2.

#### ActiveAx

The ActiveAx model (15) offers a quantitative approach to evaluate average axonal diameter within fiber bundles. This method is particularly advantageous as it is insensitive to fiber orientation. However, the reliable estimation of model parameters necessitates the use of multi-shell acquisition protocols (22). Additionally, recent studies have established correlations between water diffusivity values and axonal radius (23; 24), whereas the original ActiveAx work employed fixed diffusivity values (15). Therefore, we investigate the relationship between the minimum achievable CRLB and the intrinsic/parallel and perpendicular diffusivity values and show the results in Fig. 2 C. The identified diffusivity range aligns with previous research, which we further detail in the Discussion section. The analysis utilizes two perfectly aligned gradient directions, the first parallel to the intrinsic diffusivity and the second along the perpendicular diffusivity. To evaluate the impact of diffusivity estimation on CRLB, the axonal radius index (*R*) is varied across extreme values (0.01 and 11 *µm*), with a fixed intracellular volume fraction (*v*_*ic*_) of 0.5.

Fig. 2, Panel C, presents results for higher diffusivity regimes, characteristic of white matter. Conversely, Supplementary Fig. S3 depicts results for lower diffusivity regimes, commonly observed in gray matter or the developing brain. The two sub-figures in Panel C correspond to distinct diffusion-weighting parameter sets: low weighting with Δ = 10 ms, *δ* = 5 ms, and high weighting with Δ = 75 ms, *δ* = 65 ms. Analysis reveals an inverse proportionality (approximately reciprocal relationship) between optimal b-values (at the minimum CRLB) and diffusivity values. This trend persists regardless of the chosen diffusion-weighting parameters (Δ and *δ*) or the axonal radius index (*R*). Interestingly, the *R* value significantly influences signal attenuation only for the perpendicular diffusivity component, not the parallel component (Supplementary Fig. S3). This behavior may be attributed to the inherent diffusion properties within cylindrical compartments like axons. Parallel diffusivity within these compartments exhibits minimal dependence on the cylinder’s radius. Conversely, a larger cylinder radius is likely to elevate perpendicular diffusivity values, potentially leading to more isotropic diffusion.

#### Ball-and-Three-Sticks

The Ball-and-Stick model decomposes water diffusion into two distinct compartments: an isotropic compartment, which signifies unrestricted water diffusion, commonly referred to as the “ball,” and restricted diffusion compartments, representing diffusion constrained along specific fiber orientations, known as “sticks.” These sticks correspond to perfectly linear diffusion compartments, suitable for modeling white matter fiber tracts (16). Here, we employ three stick compartments to model three distinct fiber tracts within a voxel and incorporate a ball to capture the isotropic diffusion. Additionally, we use three gradient pulses precisely aligned with the three stick compartments. Each stick is assigned characteristic diffusivity values (*λ*_1_ = 1.74 x 10^−3^, *λ*_2_ = 1.0 x 10^−3^, and *λ*_3_ = 0.5 x 10^−3^ *mm*^2^.*s*^−1^), all falling within a biologically relevant range. Fig. 2, Panel D represents results obtained using two pairs of extreme PGSE parameters: Δ = 75 ms, *δ* = 65 ms (high weighting) and Δ = 10 ms, *δ* = 5 ms (low weighting). The optimal b-value (at minimum CRLB) for achieving the lowest variance in diffusivity estimates is approximately the reciprocal of the corresponding diffusivity value. This observation conforms to the findings from other biophysical models, such as the diffusion tensor, NODDI, and ActiveAx models.

### Optimal scanning protocols for comprehensive estimation of parameters in biophysical models

#### Diffusion Tensor

Fig. 3 explains the impact of diffusion gradient directions and signal-to-noise ratio (SNR) on the accurate estimation of all six parameters for the diffusion tensor model. To simulate a diverse array of axonal fiber configurations, orientation angles were uniformly sampled within the range of [0.01, *π*] (rad). We used ten samples for each of the three orientation angles, resulting in a total of 1000 unique diffusion tensor orientations.

**Figure 3.**
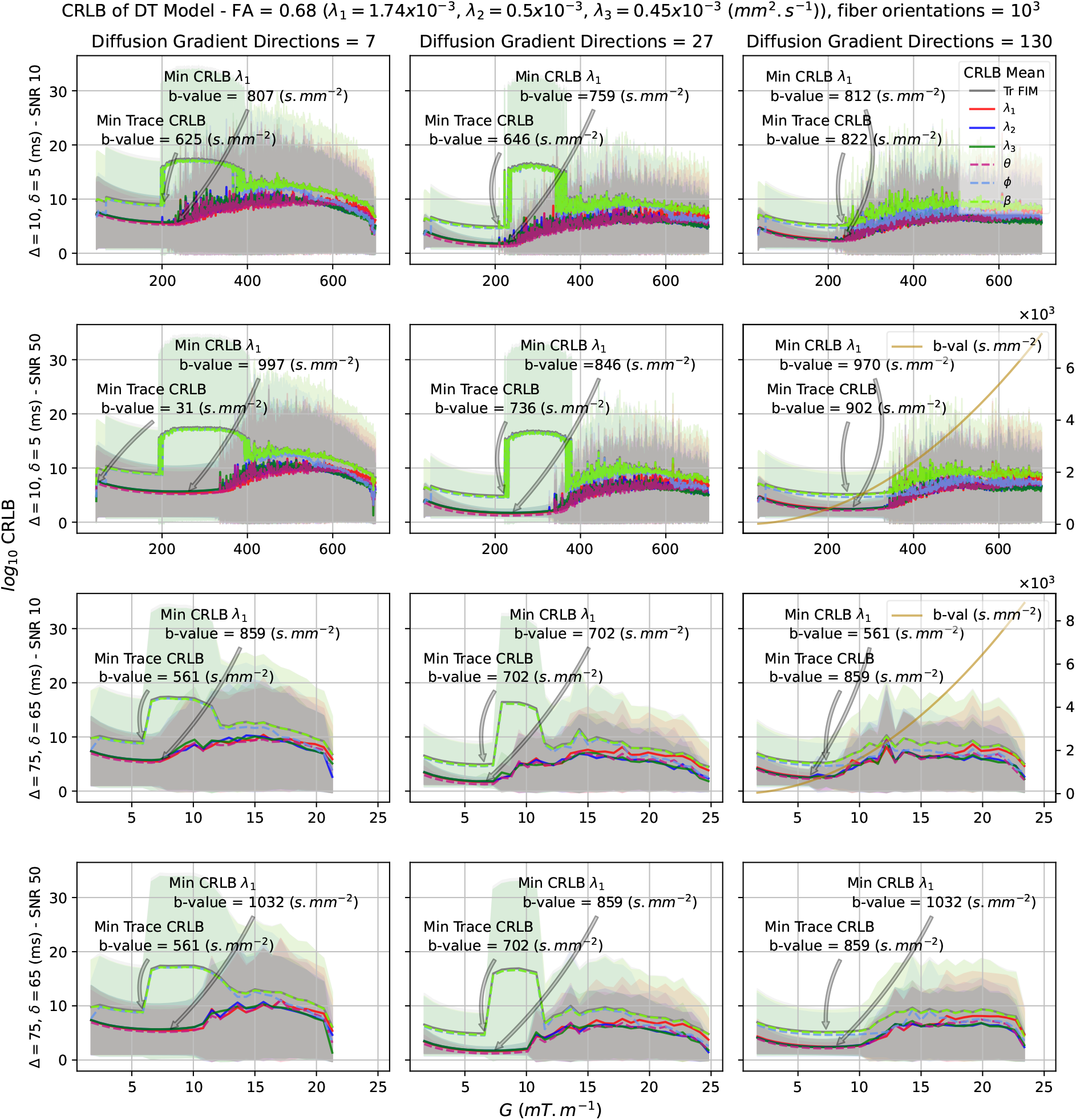
CRLB for the diffusion tensor (DT) model parameters. CRLB variability is shown for different numbers of gradient directions and b-values, using 1000 unique diffusion tensor orientations at SNR values of 10 and 50. Diffusivities were constrained within the human brain’s feasible range (illustrated by FA = 0.68). The sub-figures share a common y-axis, showing log-scale CRLB plots for all six diffusion tensor model parameters. The x-axis shows the magnitude of the MR system gradient value *G*(*mT*.*m*^−1^) with the top two rows featuring different G ranges than the bottom two rows, reflecting variations in the choice of Δ and *δ* (ms). b-values are shown in the rightmost figures of the second and third rows. Each column corresponds to three diffusion gradient tables featuring 7, 27, and 130 directions, uniformly distributed in q-space (25). Sub-figures show plots for the mean CRLB values of parameters and their sum (trace of the inverse of FIM), with the corresponding color-coded shaded areas indicating the 95 percent confidence intervals. The results from the ‘kneedle’ algorithm (26) are not shown because it would get stuck at the initial steep rise in the CRLB curve, preventing it from accurately identifying the maximum curvature point.

In a prior study focusing on SNR analysis across various brain regions, SNR values between 10 and 50 were estimated (27). Consequently, Rician noise at these two SNR levels was introduced into the simulated data. Additionally, diffusivity values were constrained within the feasible range of the human brain (17), resulting in *FA* = 0.68, distinct from the diffusivity values in Fig. 2 A. In Fig. 3, the sub-figures share a common y-axis, presenting *log*_10_CRLB plots for all six variables of the diffusion tensor model. The sub-figures also share the x-axis, displaying the magnitude of the gradient value G; however, we note that the top two rows feature different ranges for G, compared to the bottom two rows. This variation arises from the need to cover the entire space of PGSE variables. Specifically, two pairs of diffusion time and gradient duration, similar to those in Fig. 2 (i.e., Δ = 10 ms, *δ* = 5 ms for the upper two rows, and Δ = 75 ms, *δ* = 65 ms for the lower two rows), were set. For reference purposes, plots representing b-values are shown in the rightmost figures of the second and third rows, with the b-values scale indicated on the right y-axis.

Each column in Fig. 3 displays results associated with three distinct gradient tables, each featuring 7, 27, and 130 directions, respectively. The number of gradient directions was chosen to compare the results with previous works (27; 28; 29; 30) and to demonstrate how limited scanning time impacts the precision of model parameter estimation (please see Methods section for further discussion). These directions were obtained using a toolbox from INRIA, which provides uniformly distributed directions in q-space shells (29).

Each sub-figure in Fig. 3 displays plot lines representing mean values of the CRLB for all six parameters of the model, as well as the sum, i.e., diagonal elements and the trace of the CRLB matrix. The corresponding-colored shaded area for each line illustrates the 95 percent confidence interval. As expected, for a small number of gradient directions (i.e., 7), the mean CRLB exhibits higher values. Specifically, orientation angles associated with low diffusivity values, i.e., *ϕ* and *β*, show the highest mean CRLBs with a broad confidence interval. All sub-figures are annotated with the optimal b-values for the primary diffusivity and the trace of the inverse of FIM. It is evident, particularly from the right column, that an increase in pulse gradient directions leads to improved confidence intervals for all parameters, with a particular impact on orientation angles associated with low diffusivity values *ϕ* and *β* (rad), which pose the greatest challenge in terms of estimation. Additionally, the results indicate that with a low SNR of 10, the noise floor is reached earlier. For instance, in the top-right sub-figure, the minimum trace of the CRLB is observed at a b-value of 822 (*s*.*mm*^−2^). Conversely, when maintaining the same configuration but with an SNR of 50 (second row, rightmost column), the optimal CRLB value is achieved at a higher b-value of 902 (*s*.*mm*^−2^). To accurately estimate lower diffusivities and their corresponding orientation angles, employing a larger number of gradient directions and achieving higher SNR is important. In a broad sense, b-values ranging from 900 to 1000 (*s*.*mm*^−2^) are considered optimal to estimate all six parameters of the diffusion tensor, which is already the standard for clinical diffusion-weighted imaging (19; 28; 31).

The summary of results for optimal optimal experiment design for diffusion tensor model is as follows.

- The diffusivity significantly influences the selection of the optimal b-value, displaying an inverse relationship. Lower diffusivities require higher b values, increased SNR, and a greater number of gradient directions.
- More gradient directions improve the accuracy of parameter estimation by reducing CRLB and minimizing the confidence interval. However, a low SNR can compromise accuracy, even with a substantial number of directions, reaching the noise floor earlier.
- Optimal scanning protocols vary based on tissue properties and research objectives. For example, in white matter where anisotropic diffusion dominates, higher b-values may be required for accurate estimation of tissue properties.
- For the diffusion tensor model, achieving the minimum CRLB is determined by b-values, theoretically irrespective of the specific choices for *G* (*mT*.*m*^−1^), Δ, and *δ* (ms). However, practical parameter selection is limited by hardware constraints.

#### NODDI

Building upon the scanning protocols proposed by Zhang et al. (14) for the NODDI model, we evaluate their performance using experimental design optimization. Fig. 4 presents a comparison of these protocols, replicating the experimental settings from the original study. To maintain consistency with the original study for the experiment, we fixed the model parameters as follows: parallel diffusivity (*λ*_‖_) at 1.7 x 10^−3^ *mm*^2^.*s*^−1^, perpendicular diffusivity (*λ*_⊥_) at 1.2, 0.9, 0.5 x 10^−3^ *mm*^2^.*s*^−1^, separation time (Δ) = 37.8 ms, duration times (*δ*) = 17.5 ms, and orientation dispersion (*OD*) at 0.7, 0.295, 0.079, 0.0198. We simulated 900 distinct tissue configurations, consisting of 36 unique microstructural combinations, each paired with 25 fiber orientations uniformly distributed over the half-sphere. The naming convention for the protocols follows Table 1 of Zhang et al. (14) and is summarized in Table 1 in this study. The results conform to the observations by Zhang et al. (14), demonstrating the superiority of two-shell protocols over single-shell protocols. Specifically, protocol P14 has the lowest CRLB, mirroring the empirical findings by Zhang et al. (14). Using our methods, we outline three optimized two-shell protocols for the NODDI model (also detailed in Table 1). The proposed protocols achieve a lower CRLB compared to all previous NODDI protocols. However, we systematically derives these protocols through experimental design optimization, unlike the optimal NODDI protocol P14, which was identified through empirical methods. This approach offers advantages in terms of ease of implementation and flexibility. Additionally, our method can readily adapt to any scanning constraints (e.g., time) and enables the exploration of a broader range of model parameters, as described below.

**Table 1.** Imaging Protocols for the NODDI Model: This table lists the imaging protocols used for evaluation, including those from the original NODDI paper (14) (labeled P1 to P24) as well as the new protocols proposed in our study. Our proposed protocols consistently demonstrate lower CRLB values compared to the empirically selected protocol (P14) from the NODDI publication, as shown in Fig. 4 .

| Measurements per shell | G ($mT.m^{-1}$) | b ($s.mm^{-2}$) | $log_{10}$(CRLB mean) |
| --- | --- | --- | --- |
| Single Shell Protocols in the Original NODDI Study (14) |  |  |  |
| P1 |  |  |  |
| 30 | 31.9 | 711 | 9.53 |
| P2 |  |  |  |
| 30 | 37.8 | 1000 | 10.57 |
| P3 |  |  |  |
| 60 | 53.4 | 2000 | 11.34 |
| P4 |  |  |  |
| 60 | 63.8 | 2855 | 11.41 |
| Two Shell Protocols in the Original NODDI Study (14) |  |  |  |
| P13 |  |  |  |
| 30 | 31.9 | 711 | 3.05 |
| 60 | 53.4 | 2000 |  |
| P14 |  |  |  |
| 30 | 31.9 | 711 | 2.695 |
| 60 | 63.8 | 2855 |  |
| P23 |  |  |  |
| 30 | 37.8 | 1000 | 3.19 |
| 60 | 53.4 | 2000 |  |
| P24 |  |  |  |
| 30 | 37.8 | 1000 | 2.76 |
| 60 | 63.8 | 2855 |  |
| Suggested Two Shell Optimized Protocols |  |  |  |
| SP_06_29 |  |  |  |
| 30 | 30 | 631 | 2.67 |
| 60 | 65 | 2962 |  |
| SP_07_29 |  |  |  |
| 30 | 31.84 | 711 | 2.67 |
| 60 | 65 | 2962 |  |
| SP_08_29 |  |  |  |
| 30 | 33.68 | 795 | 2.675 |
| 60 | 65 | 2962 |  |

**Figure 4.**
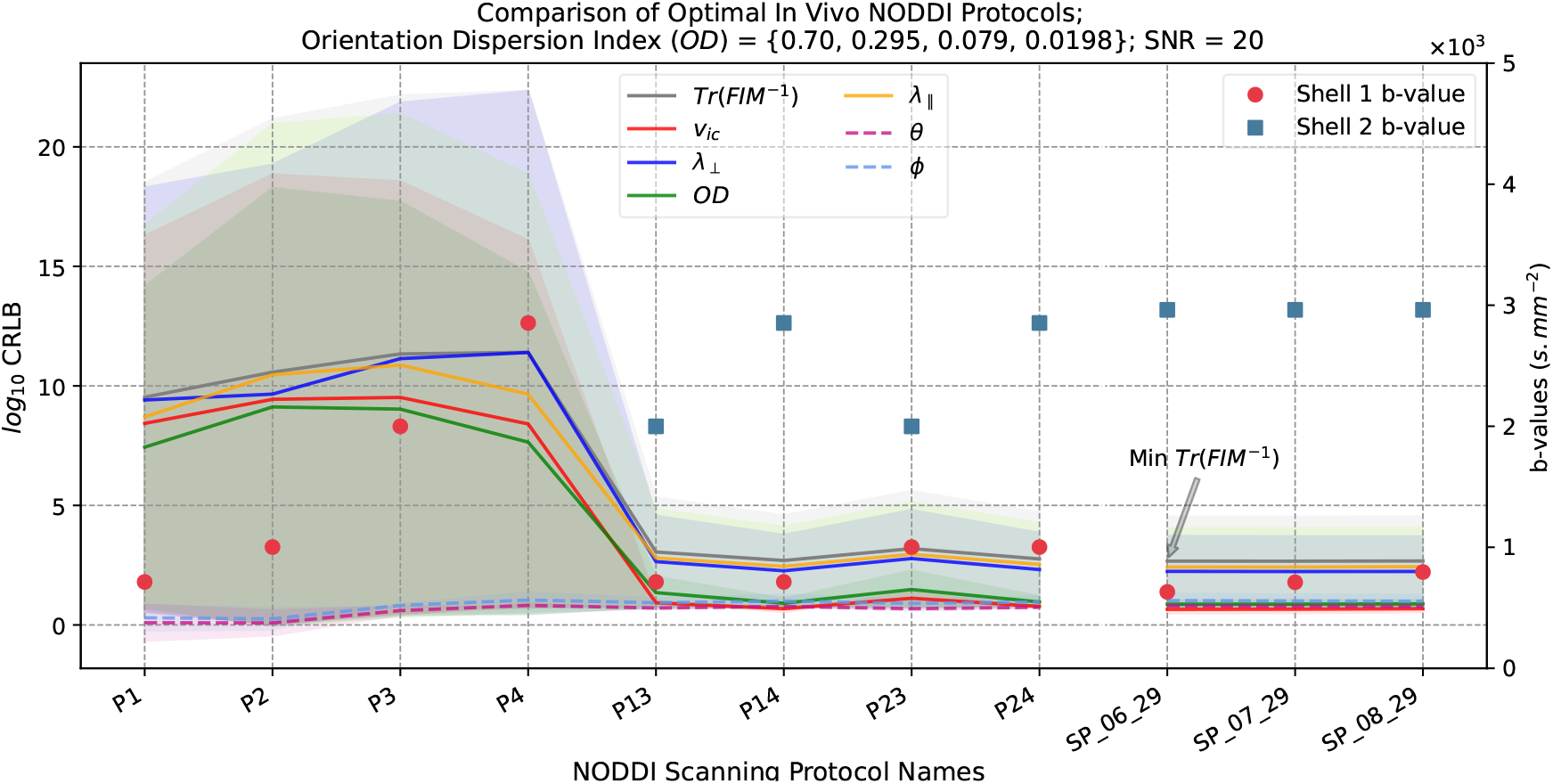
Comparison of proposed optimal NODDI protocols with those proposed in the original NODDI publication (14). The figure presents the CRLB analysis for the NODDI model parameters under various imaging protocols. The x-axis represents eight distinct scanning protocols (P1-P24) described in the original NODDI publication by Zhang et al. (14). Details of these protocols are provided in Table 1, replicated from Table 1 of the reference. The y-axis displays the mean CRLB values (in log scale) for each NODDI model parameter (*v*_*ic*_, *λ*_⊥_ and *OD*, as well as their sum i.e., trace of the inverse FIM). The corresponding color-coded shaded areas represent the 95% confidence intervals. The analysis confirms the findings of the original NODDI publication, indicating that protocol P14 exhibits the optimal performance among all the protocols considered in the original study. However, our method identified better protocols than P14, while adhering to the original study’s constraints used on model and scanning parameters. In general, protocols with a larger difference between b-values of the two shells have lower CRLB. To maintain consistency with the original study, we fixed the model parameters as follows: *λ*_‖_ = 1.7 x 10^−3^ *mm*^2^.*s*^−1^, *λ*_⊥_ = 1.2, 0.9, 0.5 x 10^−3^ *mm*^2^.*s*^−1^, Δ = 37.8 ms, *δ* = 17.5 ms, and *OD* = 0.7, 0.295, 0.079, 0.0198. We simulated 900 distinct tissue configurations, consisting of 36 unique microstructural combinations, each paired with 25 fiber orientations uniformly distributed over the half-sphere.

We extend the methodology outlined above to identify optimal scanning protocols for the NODDI model in a more general context, that is without prior assumptions on model parameters. Fig. 5 shows the dependency of the CRLB for the NODDI parameters on acquisition parameters such as *G* (*mT*.*m*^−1^), Δ, *δ* (ms), and the number of gradient directions. The x-axis of the figure represents the gradient strength, *G* (*mT*.*m*^−1^), with two columns corresponding to different ranges of *G* values determined by the choice of diffusion time and diffusion weighting factor, as indicated above each column. The corresponding b-value distributions for these ranges are depicted in the middle row. The y-axis displays the log scaled CRLB for all six NODDI parameters. Each plot shows different colored lines representing the CRLB for each model parameter, with shaded areas indicating the corresponding 95% confidence intervals. The rows of plots display results for different numbers of gradient directions: 30, 60, and 90. The NODDI model parameters were constrained to established human brain ranges as defined in the original NODDI study (14), with 36 distinct tissue configurations generated using the Cartesian product of extreme parameter values to maximize variability in the results (see methods section). Unlike previous analyses, diffusivity values are not fixed, showcasing that CRLB for model parameters decreases with increasing scanning time (or number of gradient directions). However, optimal b-values in all cases remain approximately 2200 *s*.*mm*^−2^, as indicated by the gray arrow, while the point of maximum curvature varies between 900 and 1300 *s*.*mm*^−2^. The points of maximum curvature are marked with red arrows, identified by the “kneedle” (26) algorithm with default settings. Although these are not the absolute optimal points, they represent the optimal trade-off between *G* / b-value and CRLB. Beyond these points, further increase in *G* brings minimal improvement in CRLB.

**Figure 5.**
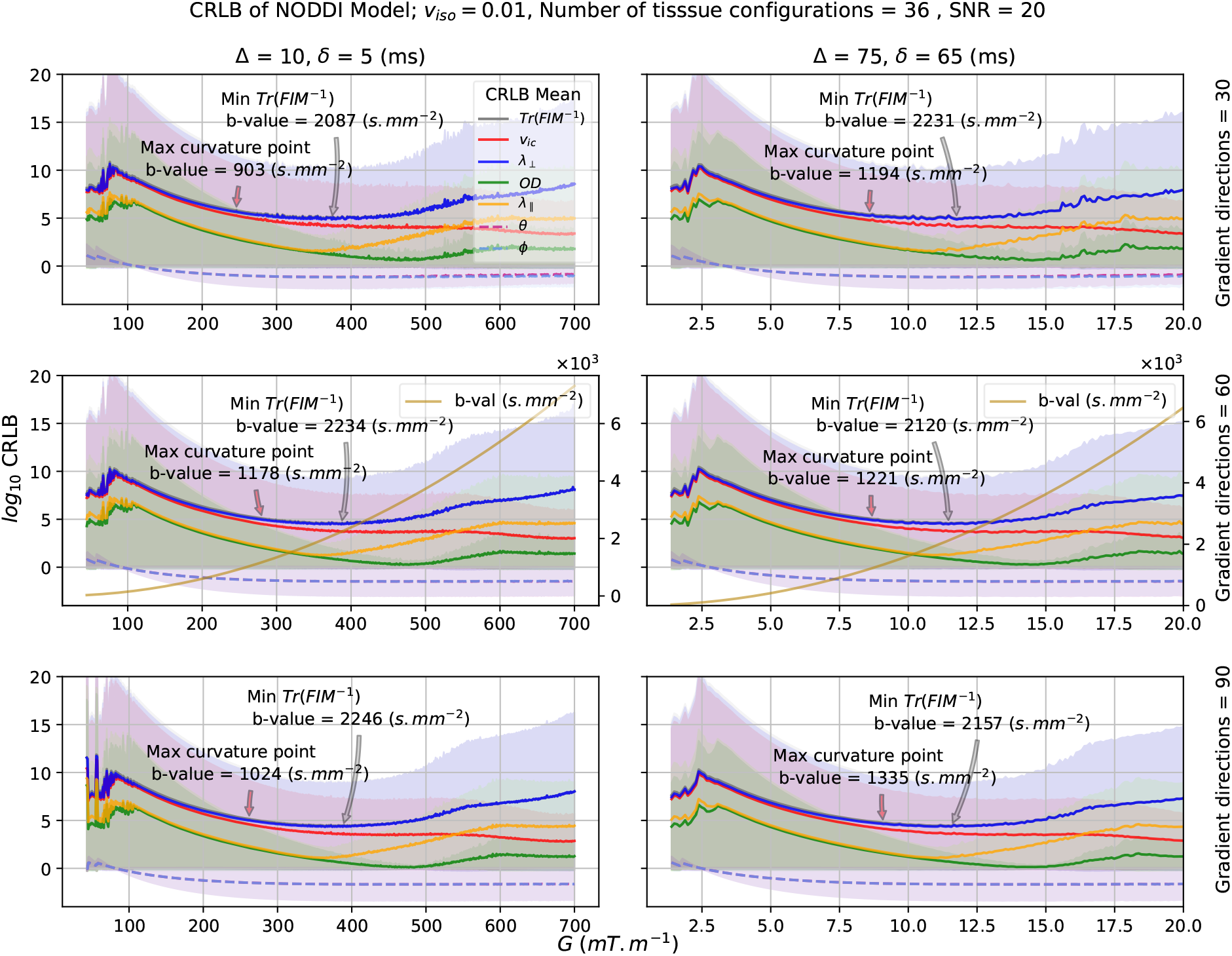
CRLB for NODDI model parameters. CRLB variability is shown for multiple gradient directions and b-values across 36 unique tissue configurations at SNR 20. NODDI parameters were constrained within the human brain’s feasible range and consistent with the original NODDI study. The sub-figures share a common y-axis, displaying *log*_10_CRLB plots for all six parameters of the NODDI model. The x-axis represents the magnitude of the MR system gradient value *G* (*mT*.*m*^−1^) with the two columns featuring different separation, Δ (ms), and duration times, *δ* (ms) of the diffusion encoding gradients. Sub-figures showcase plots for log-scaled mean CRLB values of model parameters and their sum (i.e., *Tr*(*FIM*^−1^)), with the corresponding-color-coded shaded areas indicating the 95% confidence interval. Each row corresponds to results obtained with 30, 60, and 90 diffusion gradient directions. The middle row displays b-values plots, highlighting optimal b-values corresponding to the minimum CRLB for each configuration. While the minimum CRLB indicates the point of lowest uncertainty, the maximum curvature points, detected by the ‘kneedle’ algorithm (26), highlights an optimal trade-off: the point where further increasing G or b-values produces diminishing improvements in reducing the CRLB. Although these maximum curvature points are not the absolute minimum in terms of uncertainty, they are the most informative because they mark the ‘sweet spots’ where investing in higher G or b-values no longer results in significant gains

#### ActiveAx

Fig. 6 shows the relationship between the CRLB of ActiveAx model parameters and the acquisition parameters, similar to the experiment configuration detailed in Fig. 5 for the NODDI model. Here, we employ 48 distinct tissue microstructural configurations generated through the Cartesian product of extreme model parameter values to ensure results variability. The key finding from Fig. 6 is that minimizing CRLB in the ActiveAx model relies not only on b-values but also critically on the choice of *G* (*mT*.*m*^−1^), Δ, and *δ* (ms). The figure highlights that the parameter *R* (axon radius index measured in *µm*) consistently exhibits the highest CRLB, indicating that it is the most challenging parameter to estimate accurately. Moreover, it suggests the presence of potential optimal combinations of *G*, Δ, and *δ* values within the unexplored ranges depicted in the two columns of the figure.

**Figure 6.**
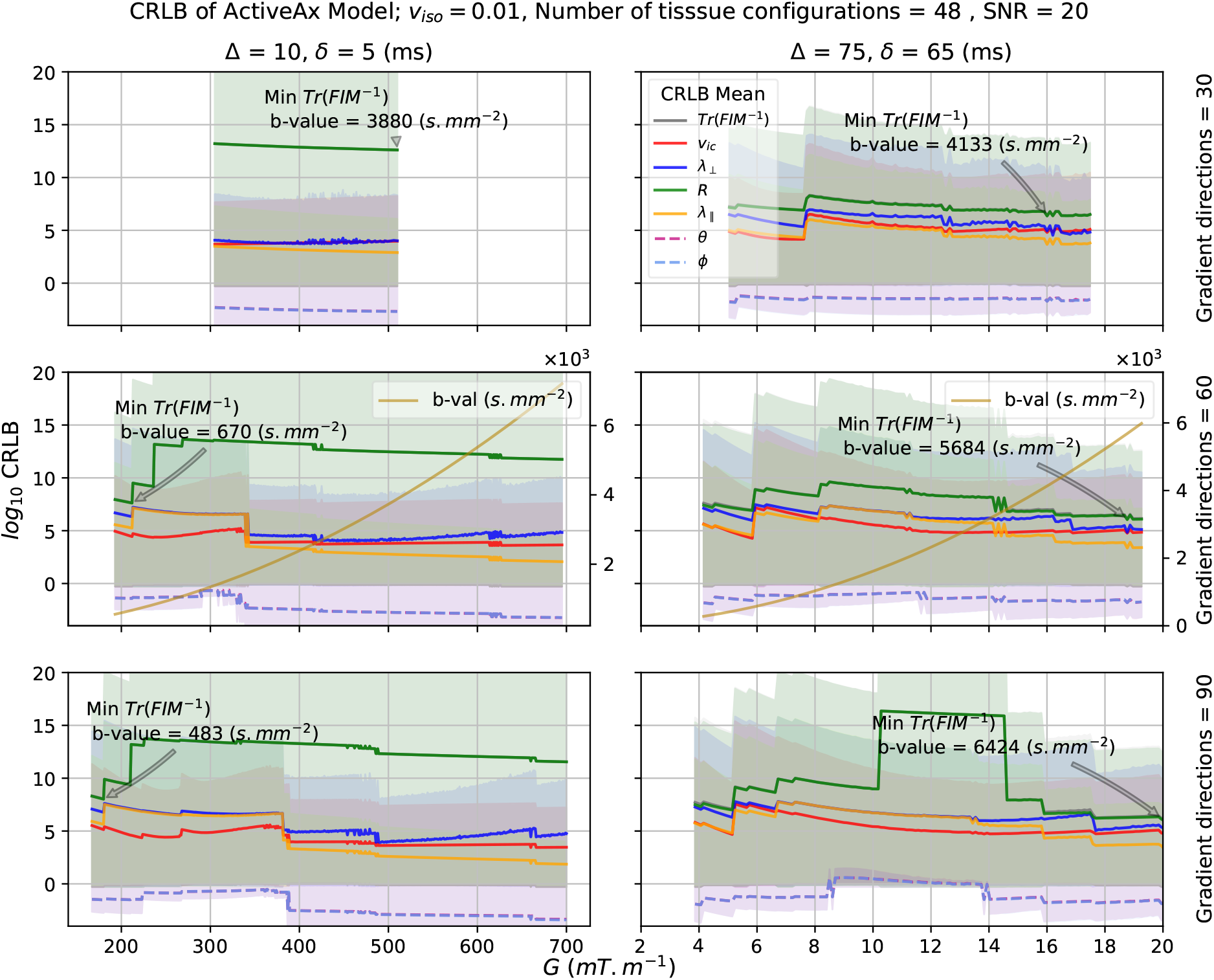
CRLB for ActiveAx model parameters. CRLB for ActiveAx model parameters. CRLB analysis of ActiveAx model using 48 unique tissue configurations at SNR = 20 is shown. ActiveAx parameters were constrained within the human brain’s feasible range. The sub-figures share a common y-axis, displaying *log*_10_CRLB plots for all six parameters of the ActiveAx model. The x-axis represents the magnitude of the MR system gradient value *G* (*mT*.*m*^−1^) with the two columns featuring different separation time, Δ and duration time *δ* (ms) of the diffusion encoding gradients. Sub-figures showcase plots for log-scaled mean CRLB values of the model parameters and their sum (i.e., *Tr*(*FIM*^−1^)), with the corresponding color-coded shaded areas indicating the 95% confidence interval. Each row shows results for 30, 60 and 90 gradient directions. The middle row shows b-values plots. The optimal b-values corresponding to the minimum CRLB are shown for each configuration. The results from the ‘kneedle’ algorithm (26) are not shown because it would get stuck at the initial steep rise in the CRLB curve, preventing it from accurately identifying the maximum curvature point.

To assess the efficacy of our proposed method for optimizing multi-shell protocols for the ActiveAx model, we compared our protocols with those presented in prior studies by Dyrby et al. (32) and Alexander et al. (15). These studies provide protocols for ex vivo and in vivo cases, which we examine in Fig. 7. Our approach demonstrates improvements for both 3-shell and 4-shell protocols in each scenario.

**Figure 7.**
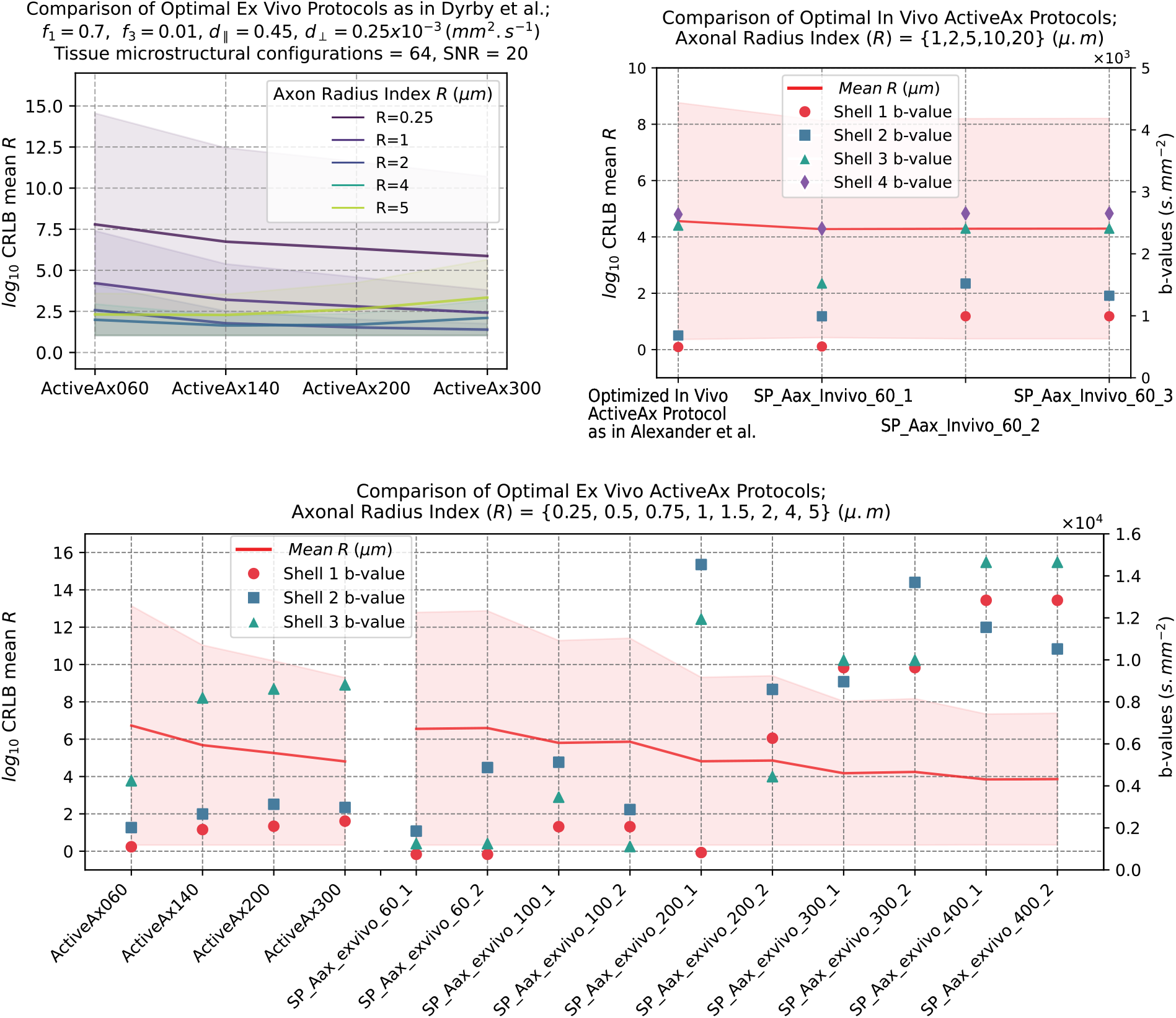
Comparison of proposed optimal ActiveAx protocols with those from prior literature. The figure compares protocols identified in our work with those from the original ActiveAx study (15) and Dyrby et al. (32) studies. The x-axis represents protocol names, following the original naming convention (details in Table 2). The left y-axis displays the log-scaled mean CRLB values of the axonal radius index *R*, with shaded areas representing the 95% confidence interval. The right y-axis shows b-values. To align with the original work, we fixed the following model parameters: for in vivo, parallel diffusivity (*λ*_‖_) to 1.7 × 10_−3_ *mm*_2_.*s*_−1_ and perpendicular diffusivity (*λ*_⊥_) to 1.2 × 10^−3^ *mm*^2^.*s*^−1^; for ex vivo, *λ*_‖_ = 0.45 × 10^−3^ *mm*^2^.*s*^−1^ and *λ*_⊥_= 0.25 × 10^−3^ *mm*^2^.*s*^−1^. Intracellular volume fraction (*f*_1_) = 0.7, isotropic volume fraction (*f*_3_) = 0.01, and SNR = 20. Top left: Comparison of optimal 3-shell ex vivo protocols by Dyrby et al (32). Protocols with higher *G*_*max*_ exhibit increased CRLB values for larger *R*, while protocols with lower *G*_*max*_ are less effective for estimating smaller *R* values, in agreement with Fig. 1 of Dyrby et al. (32). Bottom: Comparison of protocols proposed by Dyrby et al. (32) with those optimized by our method. Our protocols achieve lower CRLB values, particularly when *G*_*max*_ = 60, even with lower b-values across all shells. For higher *G*_*max*_, improved CRLB require higher b-values (see Table 2). Top right: Comparison of optimal 4-shell in vivo protocol by Alexander et al (15). Three suggested protocols show better CRLB with lower b-values compared to the original protocol.

Dyrby et al. (32) optimized ex vivo 3-shell protocols at various maximal gradient strengths (*G*_*max*_), with protocols named according to *G*_*max*_ values (e.g., ActiveAx060 for *G*_*max*_ = 60 *mT*.*m*^−1^). As shown in the top-left sub-figure of Fig. 7, we compare these protocols based on their ability to estimate the axonal radius index (*R*). We observe that protocols with higher *G*_*max*_ exhibit increased CRLB values (i.e., lower estimation accuracy) for larger *R* values, while protocols with lower *G*_*max*_ are less effective for estimating smaller *R* values. This finding explains Fig. 1 of Dyrby et al. (32), which reported an overestimation of large *R* values at higher *G*_*max*_. These results also suggest that multi-shell protocols incorporating multiple G values may be necessary to accurately estimate a range of *R* values within a region of interest in brain tissues.

**Table 2.** Multi-shell Optimized Imaging Protocols for the ActiveAx Model. This table presents optimized multi-shell protocols developed for the ActiveAx model, for both in vivo and ex vivo cases. For comparison, previously published optimal ActiveAx protocols, including those from the original ActiveAx study (15) and Dyrby et al. (32), are also included. The search space for scanning parameters (i.e., gradient strength *G*, diffusion time Δ and diffusion gradient duration *δ* was constrained to the ranges specified in these publications. Our results show that, even within these predefined constraints, improved protocols with lower CRLB can be found. CRLB plots for these protocols are shown .in Fig. 7

| Measurements per shell | $G(mT.m^{-1})$ | $\delta$ (ms) | $\Delta$ (ms) | $b(s.mm^{-2})$ | $log_{10}(CRLB\ mean)$ |
| --- | --- | --- | --- | --- | --- |
| Ex Vivo Protocols by Dyrby et al. (32) (3 shells) |  |  |  |  |  |
| ActiveAx060 |  |  |  |  |  |
| 89 | 60 | 15.9 | 22.4 | 1121 | 6.7329 |
| 98 | 54 | 15.9 | 43.3 | 2032 |  |
| 105 | 60 | 26.4 | 32.9 | 4312 |  |
| ActiveAx140 |  |  |  |  |  |
| 100 | 140 | 10.4 | 16.5 | 1863 | 5.6796 |
| 105 | 108 | 11.2 | 29.9 | 2765 |  |
| 84 | 140 | 17.4 | 23.9 | 7636 |  |
| ActiveAx200 |  |  |  |  |  |
| 102 | 200 | 7.7 | 14.2 | 1989 | 5.2585 |
| 106 | 153 | 8.8 | 25.5 | 2943 |  |
| 81 | 200 | 13.8 | 20.4 | 8757 |  |
| ActiveAx300 |  |  |  |  |  |
| 103 | 300 | 5.6 | 12.1 | 2081 | 4.8089 |
| 106 | 219 | 7.0 | 20.4 | 3080 |  |
| 80 | 300 | 10.5 | 16.9 | 9542 |  |
| Suggested Optimized Ex Vivo Protocols (3 shells) |  |  |  |  |  |
| SP_Aax_exvivo_60_1 ($G_{max} = 60$) | | | | | |
| 100 | 60 | 15.6 | 17 | 2081 | 6.5542 |
| 100 | 60 | 20.4 | 24 | 3080 |  |
| 100 | 60 | 10.8 | 45 | 9542 |  |
| SP_Aax_exvivo_100_1 ($G_{max} = 100$) | | | | | |
| 100 | 100 | 15.6 | 17 | 2056.36 | 5.802 |
| 100 | 100 | 20.4 | 24 | 5125.74 |  |
| 100 | 100 | 10.8 | 45 | 3457.92 |  |
| SP_Aax_exvivo_200_1 ($G_{max} = 200$) | | | | | |
| 100 | 200 | 6 | 10 | 824.93 | 4.8168 |
| 100 | 200 | 18 | 21.66 | 14538.88 |  |
| 100 | 200 | 10 | 45 | 11934.84 |  |
| SP_Aax_exvivo_300_1 ($G_{max} = 300$) | | | | | |
| 100 | 300 | 12 | 14.375 | 9628.55 | 4.1769 |
| 100 | 300 | 6 | 40.625 | 8961.51 |  |
| 100 | 300 | 6 | 45 | 9976.57 |  |
| SP_Aax_exvivo_400_1 ($G_{max} = 400$) | | | | | |
| 100 | 400 | 11.33 | 12.5 | 12835.43 | 3.8416 |
| 100 | 400 | 6 | 30 | 11549.10 |  |
| 100 | 400 | 6 | 45 | 11549.10 |  |
| In Vivo Protocol by Alexander et al. (15) (4 shells) |  |  |  |  |  |
| 90 | 58 | 12 | 80 | 2636.33 | 4.5614 |
| 90 | 46 | 15 | 77 | 2454.70 |  |
| 90 | 57 | 5 | 87 | 496.33 |  |
| 90 | 60 | 13 | 20 | 682.54 |  |
| Suggested Optimized In Vivo Protocols (4 shells) |  |  |  |  |  |
| 90 | 60 | 16.25 | 20 | 992.73 | 4.2763 |
| 90 | 60 | 20 | 30 | 2406.06 |  |
| 90 | 60 | 5 | 80 | 504.84 |  |
| 90 | 60 | 8.75 | 80 | 1521.41 |  |
| SP_Aax_invivo_60_2 ($G_{max} = 60$) | | | | | |
| 90 | 60 | 16.25 | 20 | 992.73 | 4.2914 |
| 90 | 60 | 20 | 30 | 2406.06 |  |
| 90 | 60 | 12.75 | 70 | 2651.77 |  |
| 90 | 60 | 8.75 | 80 | 1521.41 |  |
| SP_Aax_invivo_60_3 ($G_{max} = 60$) | | | | | |
| 90 | 60 | 16.25 | 20 | 992.73 | 4.2947 |
| 90 | 60 | 20 | 30 | 2406.06 |  |
| 90 | 60 | 8.75 | 70 | 1324.04 |  |
| 90 | 60 | 12.75 | 70 | 2651.7 |  |

The lower sub-figure in Fig. 7 compares the protocols proposed by Dyrby et al. (32) with those optimized by our method under similar experimental constraints on both scanning and model parameters. Our results indicate that the protocols generated by our method achieve better (lower) CRLB values, particularly when *G*_*max*_ = 60, even with lower b-values across all shells (which makes our proposed protocols more widely applicable to various MR scanners). For higher *G*_*max*_ values, however, achieving improved CRLB necessitates the use of higher b-values. Table 2 provides details for the highest-performing protocol at each G value, while Fig. 7 presents two representative protocols per G value.

For the in vivo case, Alexander et al. (15) proposed a 4-shell protocol, which we evaluate in the top-right sub-figure of Fig. 7. We applied the same scanning and model constraints as those in the original study, demonstrating that our method can identify more effective protocols within these constraints, even at reduced b-values. For simplicity, only the three best protocols identified by our method are shown in Fig. 7, with detailed information provided in Table 2.

#### Ball-and-Three-Sticks

Fig. 8 analyzes the Ball-and-Three-Sticks model. Here, we estimated all the twelve parameters associated with the three sticks, including diffusivity values, orientation angles, and volume fractions. To explore combinations of extreme yet biologically plausible parameter values, we generated 648 unique tissue configurations. The plots in Fig. 8 reveal that diffusivity values are generally more challenging to estimate than the orientation angles and volume fractions of the sticks. Increasing the gradient strength (*G*) and b-values typically leads to a decrease in CRLB. However, the optimal b-value range falls approximately between 5000 and 7000 *s*.*mm*^−2^, which presents challenges for in vivo (especially clinical) dMRI protocols. Fortunately, the points with the maximum curvature on the *Tr*(*FIM*^−1^) for all variables represent an optimal trade-off between b-value selection and achieving a low CRLB. The optimal value in this case is found to be around 3000 *s*.*mm*^−2^, suggesting that reliable estimation of the three-stick model parameters can be obtained without requiring prior knowledge or assumptions regarding the diffusivity values. A similar analysis with the Ball-and-Two-Sticks model with fixed diffusivity values is given in Supplementary Fig. S4 .

**Figure 8.**
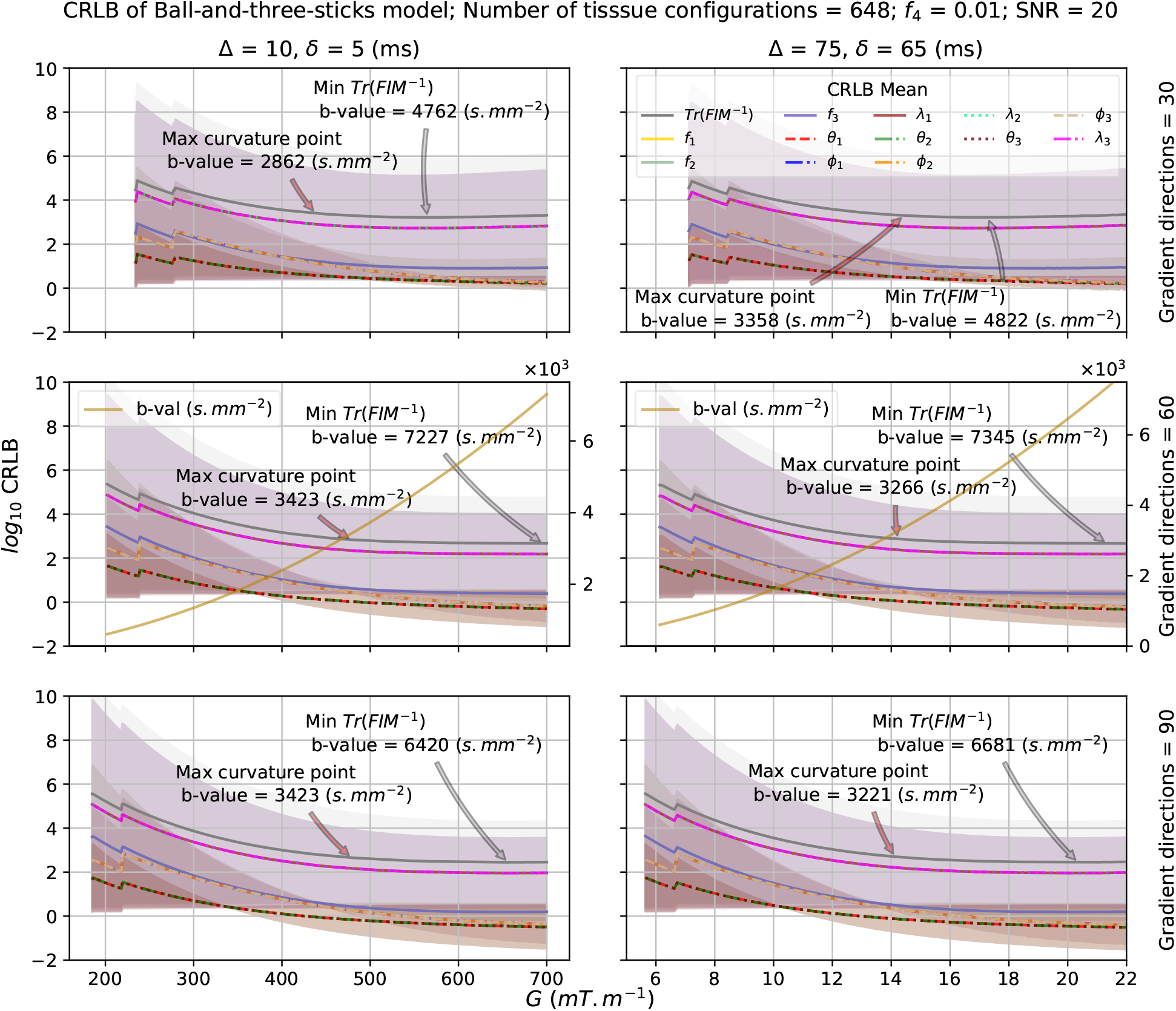
CRLB for ActiveAx model parameters. CRLB for ActiveAx model parameters. CRLB analysis of ActiveAx model using 48 unique tissue configurations at SNR = 20 is shown. ActiveAx parameters were constrained within the human brain’s feasible range. The sub-figures share a common y-axis, displaying *log*_10_CRLB plots for all six parameters of the ActiveAx model. The x-axis represents the magnitude of the MR system gradient value *G* (*mT*.*m*^−1^) with the two columns featuring different separation time, Δ and duration time *δ* (ms) of the diffusion encoding gradients. Sub-figures showcase plots for log-scaled mean CRLB values of the model parameters and their sum (i.e., *Tr*(*FIM*^−1^)), with the corresponding color-coded shaded areas indicating the 95% confidence interval. Each row shows results for 30, 60 and 90 gradient directions. The middle row shows b-values plots. The optimal b-values corresponding to the minimum CRLB are shown for each configuration. The results from the ‘kneedle’ algorithm (26) are not shown because it would get stuck at the initial steep rise in the CRLB curve, preventing it from accurately identifying the maximum curvature point.

## Discussion

### Diffusion Tensor

Fitting the diffusion tensor model to dMRI data (DTI) is widely employed to explain brain tissue microstructure due to its simplicity and reproducibility. Over the past two decades, a multitude of studies has investigated optimizing scanning protocols for DTI. These investigations have looked into the influence of factors such as the number of gradient directions, parameters of the PGSE sequence, and noise on the estimation of tissue biomarkers derived from DTI. Furthermore, research has explained the impact of specific tissue properties on the estimation of tissue microstructure using the DTI model and tailored scanning protocols. Here, we summarize key past findings from the literature and comprehensively examine how the outcomes of our study align, potentially diverge, and reconcile results from the literature.

#### Optimal b-values

The b-value in diffusion-weighted imaging is measured in *s*.*mm*^−2^, the reciprocal of diffusivity units. Higher b-values increase the degree of diffusion-weighting, causing signal intensity loss (19). Similarly, elevated diffusivity values lead to more dMRI signal loss, indicating greater diffusion. Previous studies suggest a simple rule of thumb for selecting optimal b-value i.e., b-value when multiplied by the ADC of the tissue under investigation should be close to 1 (18) or approximately 1.1 (33). Also, previous work shows that for DTI, optimal b-values range from 900 to 1000 (18; 28; 31; 34). Our results in Fig. 2A align with the previous literature, however, derived solely from CRLB for the diffusion tensor model.

#### Number of gradient directions

Theoretically, for the estimation of the six parameters of the diffusion tensor, employing at least six non-collinear diffusion encoding directions along with one minimally T2-weighted low b-image (b = 0 *s*.*mm*^−2^) can be sufficient (28; 29). However, Jones et al. (30) found that for the estimation of all parameters of the diffusion tensor, approximately 30 gradient directions serve as a practical compromise between quality and scanning time. They also observed that increasing the number of orientations beyond 30 did not lead to improved tensor orientation estimation. However, there are contrasting findings as well. Another study, conducted by Correia et al. (27), suggests that, for estimating fiber orientation, an acquisition scheme utilizing as many directions as possible and only one b-value produces the best results in terms of accuracy and stability. In our study involving 7, 27, and 130 gradient orientations, we find that a higher number of orientations improves fiber orientation estimates, consistent with the findings of Correia et al. (27).

#### Tissue properties

Our study demonstrates that the optimal b-value for DTI can be influenced by the specific tissue type and the physiological or pathological characteristics under investigation. Diffusivity values, which vary in white matter, gray matter, cerebrospinal fluid (CSF), and in diseased tissues, may require optimal b-values that deviate from the widely accepted clinical standard of b=1000 *s*.*mm*^−2^. Following are examples from previous studies that highlight optimal b-values for DTI specific to various tissue types: (1) *Discriminating High-Grade and Low-Grade Gliomas*: A higher b-value of 3000 *s*.*mm*^−2^ proves more beneficial than b = 1000 *s*.*mm*^−2^ for distinguishing between high-grade and low-grade gliomas (35); (2) *Peripheral Nerves Imaging*: The optimum tissue-specific b-value in peripheral nerves is capped at approximately 700 *s*.*mm*^−2^ (36); (3) *Multiple Sclerosis-Related Hippocampal Microstructural Alterations*: To investigate the impact of multiple sclerosis on gray matter in a mouse model, only DTI with the highest tested b-value (2700 *s*.*mm*^−2^) revealed dendritic loss. Thus, achieving biologically meaningful gray matter DTI metrics necessitates high b-values and an adequate number of gradient directions (37).

### NODDI

NODDI is one of the most widely used multi-compartment brain tissue biophysical model due to its short scan times and the availability of various implementations. The original NODDI publication (14) identified an optimal scanning protocol for the model by empirically evaluating various single-shell and multi-shell combinations. However, subsequent research has indicated that the choice of protocol significantly affects NODDI metrics (38; 39; 40). Therefore, scanning protocol design should adapt to study specific requirements (e.g., tissue type to be studies) and available scan time. Recent investigations have examined the reproducibility of NODDI metrics based on b-value and diffusion sampling schemes (38), as well as the model’s reliability across adult and pediatric brains (39). Additionally, NODDI metrics derived from high-resolution diffusion MRI in ex-vivo mouse brains were compared with subsequent histochemical analysis of myelin and neuronal density to recommend b-values for gaining new insights into brain microstructure (40). Our work introduces a systematic framework for identifying optimal NODDI scanning protocols using CRLB analysis (Fig. 5), offering flexibility for different tissue types and scan time constraints. The following subsections discuss how our approach to NODDI protocol optimization aligns with existing research, considering the underlying assumptions and limitations of each study, including our own.

#### Simplifying assumptions on tissue microstructure

NODDI simplifies the process of parameter estimation by making several assumptions about the tissue architecture and diffusion properties. These include the presence of a single dominant fiber orientation per voxel, a fixed intrinsic parallel diffusivity value, estimation of perpendicular diffusivity using a tortuosity model, negligible water exchange between compartments, and a low volume fraction of isotropic diffusion, particularly in the white matter. These simplifications enable the parameters to be efficiently and reliably estimated from the dMRI signal.

However, several studies have demonstrated the (potentially negative) impact of predefined parallel diffusivity values, commonly set to 1.74 x 10^−3^ *mm*^2^.*s*^−1^ in vivo (20; 21; 40), on NODDI parameters estimation. For instance, a recent study examined different diffusivity values and observed their influence on other estimated parameters, suggesting that in vivo diffusivity values fall within the range of 2 - 2.5 x 10^−3^ *mm*^2^.*s*^−1^ (21). Similarly, another study using ex vivo mouse brain data, varied diffusivity from 0.6 x 10^−3^ *mm*^2^.*s*^−1^ to 1.2 x 10^−3^ *mm*^2^.*s*^−1^ for optimal fitting results, validated by histochemical analysis of myelin and neuronal density. Additionally, Grussu et al. (41) used an axial diffusivity value of 1.5 x 10^−3^ *mm*^2^.*s*^−1^ in a postmortem spinal cord study, achieving optimal fitting accuracy (21).

Based on the findings from Fig. 2B (and Supplementary Fig. S2), we suggest that tissue diffusivity values could be estimated via NODDI when b-values are approximately the reciprocal of diffusivity values. However, two key conditions must be met: 1) Low *OD*, typically found in white matter, as high *OD* can impede accurate diffusivity estimation; 2) Aligning some diffusion gradient orientations with the principal directions of diffusion is crucial, and achievable by maximizing the number of gradient orientations within scan time constraints.

#### Optimal b-value

Studies have focused on optimizing NODDI scanning protocols to achieve a practical balance between scan time and reproducibility of estimated parameters (38; 39; 40). This optimization process involves finding the best combination of diffusion weighting factors (b-values) and number of diffusion gradient directions to acquire reliable data while minimizing scan time.

One approach to protocol optimization involves evaluating the empirical reproducibility of NODDI metrics based on different b-value and diffusion sampling schemes. Parvathaneni et al. (38) investigated this concept and suggested that, for estimating the intracellular volume fraction (*v*_*ic*_), a b-value of 2500 *s*.*mm*^−2^ was optimal. In contrast, for orientation dispersion (*OD*), a b-value of 2500 or 3000 *s*.*mm*^−2^ was found to be the most suitable, with the requirement for a total number of diffusion directions exceeding 128. Another study (39) used the same scanning protocol and applied a test-retest approach to assess the reproducibility and reliability of NODDI biomarkers from both pediatric and adult subjects. Beyond in vivo studies, ex vivo investigations have also contributed to protocol optimization. A study using mouse brain data concluded that a 3-shell datasets with b-values of 1000, 4000, and 8000 *s*.*mm*^−2^ produced model parameter estimates close to those obtained using a 6-shell protocol with b-values of 500, 1000, 2000, 4000, 6000, and 8000 *s*.*mm*^−2^ (40).

Our observations in Fig. 5 partially agree with the prior investigations. The figure indicates that the optimal b-value for estimating orientation dispersion (*OD*) lies around 4000 *s*.*mm*^−2^, a value higher than the 3000 *s*.*mm*^−2^ proposed by Parvathaneni et al. (38). However, it is important to note that Parvathaneni et al. (38) utilized a maximum b-value of 3000 *s*.*mm*^−2^2. In contrast, for estimating *v*_*ic*_, our findings diverge. While Parvathaneni et al. (38) suggested a b-value of 2500 *s*.*mm*^−2^ to be optimal, Fig. 5 demonstrates that increasing b-values could potentially reduce the CRLB further, thus reducing variability in parameter estimates. Additionally, in ex vivo brain studies, brain tissues exhibit lower diffusivity compared to in vivo settings, requiring higher b-values for accurately estimating model parameters.

The results presented in this study remain independent of the model fitting algorithms utilized. Nonetheless, it is essential to recognize that investigations assessing the reproducibility of NODDI metrics, as discussed above, utilize diverse model fitting techniques encompassing DMT (42), CuDIMOT (43), AMICO (44), and the NODDI MATLAB toolbox (14). This may also impact the accuracy of parameter fitting, alongside considerations such as b-values, gradient directions, and fixed diffusivity values. Subsequent research endeavors for assessing reproducibility using empirical methods should therefore attempt to standardize fitting methods for more uniform comparisons across studies.

### ActiveAx

Axonal diameter is a critical characteristic of brain white matter, directly impacting conduction speed and influencing action potential firing rates (23; 45; 46). These processes are fundamental for evaluating brain function. Consequently, the ability to measure axonal diameter accurately, non-invasively, and in vivo using dMRI data holds immense value for investigating brain diseases and healthy development.

The ActiveAx technique addresses this need by employing a biophysical model known as the Minimal Model of White Matter Diffusion (MMWMD) (15; 22). The model leverages dMRI data to estimate the axonal radius index (*R*) within a voxel, representing the main fiber bundle. The MMWMD characterizes the intra-axonal space as a compartment with restricted water diffusion confined to a set of cylinders with uniform radii. Conversely, the extra-axonal space is modeled as a compartment exhibiting hindered water displacement perpendicular to the axons. Additionally, the model can be optimized for in vivo or ex vivo applications by incorporating compartments for cerebrospinal fluid and stationary water (22) (see Methods section for details).

The ActiveAx model offers a valuable tool for estimating axonal diameter index, but it relies on certain assumptions that may limit its accuracy. One assumption is the presence of a single fiber orientation within a voxel. Additionally, the model treats diffusivities as fixed values. A recent study by Pizzolato et al. (47) demonstrated the potential of using the PGSE sequence to estimate both parallel diffusivity (*λ*_‖_) and perpendicular diffusivity (*λ*_⊥_) within a voxel and then use the estimated *λ*_⊥_ to infer axon radius index index (*R*). Furthermore, several studies have reported a proportional relationship between *λ*_⊥_ and *R*, with *λ*_⊥_ increasing monotonically as *R* increases (23; 24).

The findings in Fig. 2C support this finding. The CRLB of *λ*_⊥_ is influenced by variations in *R*. Conversely, the CRLB of *λ*_‖_ remains largely unaffected by changes in *R*. This confirms that *λ*_⊥_ and *R* are indeed interrelated and that variations in *R* can influence the accuracy of *λ*_⊥_ estimation.

Fig. 6 comprehensively explores how the CRLB of the ActiveAx model parameters is impacted by utilizing a wide range of the model parameter values. In essence, the figure examines how varying these parameters across their extreme values influences the minimum achievable variance in our estimates of the model parameters. A key takeaway is that *R* consistently exhibits the highest CRLB across all scenarios, suggesting it is the most challenging parameter to estimate accurately. In contrast to models like DT (13), NODDI (14), and Balls-and-Stick (16; 48) that rely solely on b-values (and the number of diffusion gradients) for scanning protocol optimization, the accuracy of ActiveAx estimations depends on a combination of PGSE parameters: *G* (*mT*.*m*^−1^), Δ, and *δ* (ms). Further, to identify and compare optimal multi-shell acquisition protocols for the model, we conducted a systematic evaluation against established protocols from the literature. The results, presented in Fig. 7, demonstrate that our proposed method effectively searches across a wide range of scanning parameter combinations in multi-shell settings. This approach allows us to tailor protocol optimization to the specific constraints of the scanning hardware, as well as the number of shells and measurements per shell. By doing so, we maximize the information content of the MRI signal, for a given biophysical model, within given experimental constraints.

### Ball-and-Stick

The Ball-and-Stick model provides a simplified representation of white matter microstructure, conceptualizing axons as impermeable cylinders with negligible radius and assuming isotropic diffusion in the surrounding extracellular space. Initially introduced within a Bayesian framework for parameters estimation and tractography (16; 48), this model has been widely adopted to implement probabilistic approach in tractography toolsets within prominent software packages such as FSL (49) and FreeSurfer/ TRACULA (50; 51) for tractography analysis. However, challenges emerge when dealing with voxels containing crossing fibers, as highlighted by recent studies indicating inaccuracies in fiber parameter estimation within such scenarios (52). This issue is particularly significant given the prevalence of crossing fibers, with estimates suggesting that 60 − 90% of white matter voxels exhibit such complex geometry (53). To tackle this challenge, a simplified version of the Ball-and-Stick model was previously proposed (52). Moreover, another study introduced model specific fitting techniques employing an Expectation-Maximization algorithm to enhance model parameters estimation accuracy (54). We, therefore, focus on mitigating the issue of inaccurate parameter estimation of the Ball-and-Stick model, when applied to complex fiber geometries, by identifying optimal scanning protocols specifically tailored for the ball-and-two-sticks and ball-and-three-sticks models. Fig. 8 and Supplementary Fig. S4 illustrate the relationship between b-values, the number of gradient directions, and the CRLB for both the models. It is shown that higher b-values and a greater number of gradient directions result in lower CRLB values. These findings suggest that estimating the Ball-and-Stick model in scenarios involving fiber crossings, such as those with two or three sticks, is challenging when using scanning protocols like the one originally used by Behrens et al. (48), which includes a single low b-value of 1000 *s*.*mm*^−2^ and only 60 gradient directions.

The findings suggest that, for fixed diffusivity values and two fiber orientations/ sticks, a two-shell protocol with low b-value of 700 *s*.*mm*^−2^ and high b-value ranging from 4000 to 7000 *s*.*mm*^−2^ is recommended, with at least 30 to 60 directions in each shell. Similarly, for the ball-and-three-sticks model, with variable diffusivities in each stick, lower b-value of 3000 *s*.*mm*^−2^ and higher b-value ranging from 5000 to 7000 *s*.*mm*^−2^ is recommended, along with 30 to 60 directions in each shell.

Our study has several key limitations which require consideration to accurately interpret and apply the results: 1) Biophysical Model Generality: While common and complex biophysical models (diffusion tensor, NODDI, ActiveAx, Ball-and-Three-Sticks) were implemented to demonstrate generalizability of the method, extension to other models requires a modular approach. This likely involves integration into libraries like Dmipy (55) to facilitate customization for specific hardware and user-defined models; 2) In vivo Applicability: This work focused on demonstrating optimal dMRI experiment design by differentiable programming of tissue models and efficient GPU implementation. However, detailed in vivo experiments will be necessary to customize the findings to specific hardware constraints and tissue properties in future studies; 3) CRLB Optimization Dependence: The accuracy of our CRLB-based optimization relies on the assumption that the chosen tissue model accurately reflects the underlying tissue geometry. Deviations from this assumption can lead to inaccurate optimization results. However, we anticipate that our proposed experimental design framework will facilitate the development and assessment of novel (possibly more accurate) biophysical models; 4) dMRI Parameter Estimation Challenges: Estimating model parameters from dMRI data inherently involves a non-convex, non-linear optimization problem (56). Different optimization algorithms have their own limitations, and variations in estimated parameters can arise beyond optimal dMRI scan design considerations.

In summary, our study presents a method to optimize dMRI data acquisition for specific scan time constraints and tissue properties/biophysical model. We achieve this by leveraging advances in automatic differentiation and parallel computing to enable efficient implementation of complex biophysical tissue models. This approach offers a generalizable and scalable framework applicable to any biophysical model utilizing dMRI data for microstructure imaging.

## Methods and Materials

### Experimental design optimization for dMRI acquisition using multi-compartment biophysical models

The experiment design optimization problem aims to find the optimal acquisition parameters (*α*) (further detailed below) such that the precision of the estimated model parameters (*x*) (description follows) is maximized. In other words, we seek to minimize the sum of the standard errors associated with each model parameter in *x*.

Eq. 1 minimizes the trace of the inverse Fisher Information Matrix (FIM) with respect to the scanning parameters *α*. The FIM, denoted as *F*, depends on both parameter vectors *α* and *x*. This minimization enhances the precision of estimating model parameters (*x*), as the trace of *F* represents the sum of variances (diagonal elements) – the Cramér-Rao Lower Bounds (CRLB) for *x*. Minimizing the objective function with respect to the acquisition parameters *α* therefore ensures that dMRI data obtained using these scanning parameters yields model parameter (*x*) estimates with minimal uncertainty.

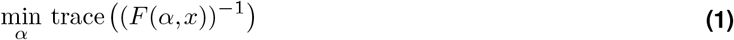

Derivation of the objective function in Eq. 1 is given in Supplementary text.

#### Acquisition Parameters (*α*) of the Pulse Gradient Spin Echo (PGSE) sequence

We aim to identify the optimal parameters for single diffusion encoding (SDE) using the Pulse Gradient Spin Echo (PGSE) sequence for biophysical modeling applications from dMRI data. The key PGSE parameters for SDE include: 1) *Gradient strength (G)*: magnetic field gradient magnitude (*m*.*T*^−1^); 2) *Diffusion pulse duration (δ)*: duration of the diffusion-sensitizing pulse (ms); and 3) *Diffusion time (*Δ): interval between the diffusion gradients (ms) (Supplementary Fig. S1B). Therefore, the set of independent parameters, denoted by *α*, is defined as:

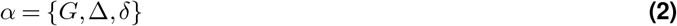

The three independent parameters collectively define the diffusion weighting factor *b* (*s*.*mm*^−2^), calculated as:

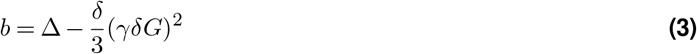

where *γ* represents the gyromagnetic ratio = 267.515 × 10^6^ *rad*.*s*^−1^.*T* ^−1^.

#### Fixed Parameters

##### Gradient Directions

A single unit vector, *ĝ* = [*g*_*x*_; *g*_*y*_; *g*_*z*_], defines a spatial direction in 3D for the PGSE sequence. Depending on available scanning time, we can employ a set of *L* such gradient directions. These are conveniently represented as a matrix, *g* ∈ℝ^*L*×3^, where each row is a unit vector like *ĝ*. In this study, we leverage a toolbox from INRIA (25) to acquire a set of *L* directions uniformly distributed across *q*-space shells.

##### Gyromagnetic Ratio (*γ*)

This constant relates magnetic field strength to nuclear precession frequency with a value of approximately 267.515 × 10^6^ *rad*.*s*^−1^.*T* ^−1^.

##### Common parameters of the biophysical tissue model functions (x)

This section details the common parameters used in the biophysical model functions representing the various tissue compartments. Model-specific functions and additional parameters will be introduced in subsequent sections.

##### Volume Fraction (f_i_)

The volume fraction of the *i*^*th*^ compartment, where *i* = 1, 2,…, *N* . These fractions sum to unity across all compartments: 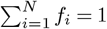.

##### Fiber orientation angles

The variables *θ*_*i*_ and *ϕ*_*i*_ respectively denote the fiber orientation angle in the *x*-*y* plane and about the *z*-axis (in radians) for the *i*^*th*^ fiber orientation.

##### Fiber orientation vector (n_i_)

Defined by *θ*_*i*_ and *ϕ*_*i*_ using the following equation:

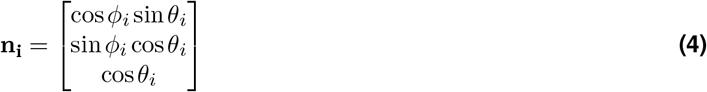

##### Water diffusivity values (λ_i_)

*Parallel diffusivity*, denoted as *λ*_‖_, signifies the diffusivity along the direction parallel to the fiber orientation vector **n**_**i**_, commonly referred to as intrinsic free diffusivity. In the literature, this parameter is frequently constrained to specific values, such as 0.6 × 10^−3^ mm^2^ · sec^−1^ for ex-vivo and 1.7 × 10^−3^ mm^2^ · sec^−1^ for in-vivo brain tissues (57). Conversely, *perpendicular diffusivity*, denoted as *λ*_⊥_, characterizes the diffusion perpendicular to **n**_**i**_. It is derived from the parallel diffusivity using a tortuosity model (58). *Isotropic diffusivity, λ*_iso_, describes the unrestricted diffusion of water molecules, and is fixed to 2.0 × 10^−3^ mm^2^ · sec^−1^ for ex-vivo and 3.0 × 10^−3^ mm^2^ ·sec^−1^ for in-vivo brain tissues. However, recent studies have reported higher diffusivity values. For instance, Howard et al. (21) reported 2 – 2.5 × 10^3^*mm*^2^.*s*^−1^ in vivo and observed that the value influences other estimation of other model parameters. Therefore, in this study, we do not fix the diffusivity values except for isotropic diffusivity and instead provide the CRLB for all diffusivity values in the models studied.

##### Details on biophysical models’ parameters

In addition to the common model parameters outlined above, this section elaborates on model-specific parameters required for a comprehensive definition of the parameter set x in the optimization problem given in Eq. 1. Below, we provide a technical overview of the commonly utilized biophysical models for brain microstructure imaging.

#### Diffusion Tensor

Normalized estimated diffusion MRI signal *ŷ*_DT_ with diffusion tensor model:

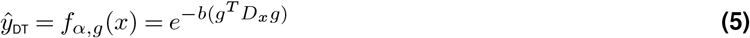

Where 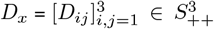 is the diffusion tensor with dependent model parameters *x* = {*λ*_1_, *λ*_2_, *λ*_3_, *θ, ϕ, β*}, where *λ*_*i*_ are the eigen values of the diffusion tensor and *θ, ϕ, β* give rotation of DT around x, y, and z-axis respectively (Supplementary Fig. S1A).

#### NODDI

The estimated dMRI signal *ŷ*_NODDI_ comprises the normalized signals from the following three compartments:

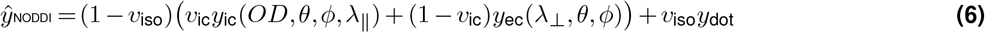

Here, *y*_ic_ and *y*_ec_ denote the normalized signals from the intra-cellular and extra-cellular compartments of NODDI model, respectively, as outlined by Zhang et al. (14). The isotropic dMRI signal function *y*_dot_ can be found in supplementary information of Farooq et al. (56). The NODDI model is parameterized by a set of dependent variables, *x* = {*θ, ϕ, λ*,_‖_ *λ*_⊥_, *v*_*ic*_, *v*_*iso*_, *OD* }, wherein *v*_*ic*_ signifies the intra-cellular volume fraction, *v*_*iso*_ represents the isotropic volume fraction, and *OD* denotes the neurite orientation dispersion index (14).

#### ActiveAx

The estimated dMRI normalized signal using ActiveAx model (15) *ŷ*_ActiveAx_ is assumed to originate from the following three compartments:

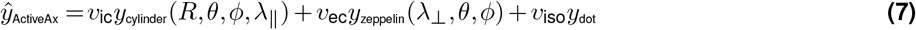

where, *v*_*ic*_, *v*_*ec*_, and *v*_*iso*_ are the intra-, extra-and isotropic volume fractions such that *v*_*ic*_ + *v*_*ec*_ + *v*_*iso*_ = 1. Details about the model functions for *y*_*cylinder*_, *y*_*zeppelin*_, and *y*_*dot*_ can be found in (15; 57) and also in the supplementary information of (56). The ActiveAx model has model parameters set *x* = {*θ, ϕ, λ*_‖_, *λ*_⊥_, *v*_ic_, *v*_ec_, *v*_ec_, *R* }, where *R* denotes axon radius index (*µm*).

##### Ball-and-Stick

The normalized estimated dMRI signal from Ball-and-N-sticks can be expressed as follows:

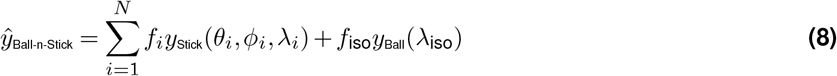

Here, 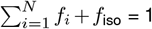, while *λ*_*i*_ is the diffusivity value along stick *i*. Model functions for *y*_Stick_ and *y*_Ball_ can be found in Behrens et al. (16) and also in supplementary information of (56). For Ball-and-Three-Sticks, the model parameters set is *x* = {*θ*_*i*_, *ϕ*_*i*_, *λ*_*i*_, *f*_*i*_ | *i* = 1, 2, 3} ∪ {*f*_*iso*_}.

### Comprehensive assessment of tissue configurations

To systematically explore the parameter space and devise an optimal scanning protocol, we establish sets containing the extreme permissible values for each model parameter. For instance, let *F* = {0.1, 0.79} denote the range of volume fractions, Θ = {0, *π/*2} encompass the allowable extreme values of *θ*_*i*_, Φ = {0, *π*} represents the minimum and maximum possible values for *ϕ*_*i*_, and *D* = {0.5, 1.74} represent the range of diffusivity values. To generate all possible combinations of parameter values across these sets, we employ the Cartesian product operation. The Cartesian product *F* × Θ × Φ ×… ×*D* yields all possible combinations of parameter values. This product is repeated *N* times, where *N* represents the number of compartments, such as the number of sticks in the Ball-and-Sticks model.

This process results in the comprehensive set of all possible combinations of model parameters. Subsequently, infeasible combinations are filtered out. For instance, combinations where the sum of volume fractions does not equal one, or where the fiber orientation within a compartment is repeated or identical, are eliminated. This rigorous approach ensures that only physically permissible parameter combinations are considered in the experiments, facilitating a thorough exploration of the parameter space.

### Differentiable programming for biophysical tissue models using dMRI

The power of biophysical models comes at the cost of complexity. Indeed, these models rely on intricate mathematical formulations to describe the diffusion process within specific tissue geometries (e.g. cylinder). Therefore, analyzing these models and optimizing MRI sequences based on them necessitates the calculation of derivatives (including CRLB) with respect to specific (and ideally, as proposed here, all) model parameters.

Manual derivations of these gradients prove challenging due to the inherent complexity of the models. The intricate nature of mathematical expressions translates into a time-consuming and laborious differentiation process, inherently prone to errors. While analytical solutions exist for simpler models (e.g., design optimization for spin echo sequences by Zhao et al. (59), they become impractical for more intricate biophysical models.

Differentiable programming offers a solution by treating the entire model as a differentiable entity. This concept unlocks powerful optimization and analysis techniques through automatic differentiation. Automatic differentiation leverages the chain rule to efficiently compute the derivatives of complex functions, ensuring both accuracy by minimizing errors through automation and efficiency by significantly reducing computation time compared to calculating analytical derivatives. The technique has been widely used in the field of deep learning in the process of backpropagation (60), physics (61), and biology (62).

In quantitative MR sequence optimization, Lee et al. (7) employed the Python library Autograd (63) for the automatic differentiation of Bloch simulations. However, Autograd is no longer under active development, and it tends to be too slow for medium to large-scale experiments as it does not implement Graphics Processing Units (GPU) support. In this work, we implemented biophysical models into the TensorFlow (12) framework to capitalize on its capabilities for automatic differentiation and GPU acceleration. It enables the exploitation of automatic differentiation benefits in computing the CRLB for biophysical models’ parameters, while also harnessing GPU computational power for accelerated derivative computations.

### Gaussian noise model assumption for minimizing CRLB

For MRI data with SNR greater than 6, the Rician noise distribution approximates the normal distribution (64). Also, the objective function for minimizing CRLB with Rician noise model involves an integral part without close form solution, which requires either a numerical solution or interpolation of values from pre-computed lookup table (2). Therefore, to compute derivatives of the biophysical model functions, a Gaussian noise model was assumed.

### Rank constancy of FIM

A full-rank FIM is essential, indicating that the information content is sufficient for parameter estimation without redundancy. Three key factors contribute to achieving a full-rank FIM:

1. *Choice of Model Parameters*: Selecting a diverse set of parameters that comprehensively capture various aspects of the tissue microstructure is crucial. This heterogeneity helps ensure the FIM is full rank and avoids parameter redundancy.
2. *Diffusion Gradient Directions*: In diffusion MRI experiments, applying magnetic field gradients along multiple directions allows for probing the tissue microstructure from various angles. This multi-angular scanning scheme enriches the information content, contributing to a full-rank FIM.
3. *Measurement Noise*: While ideally minimized, a small amount of noise in the data can positively impact the FIM’s rank. Noise can disrupt potential symmetries or degeneracies within the data that might otherwise lead to a non-full-rank FIM. Therefore, to ensure analyses are based on data with sufficient information for accurate parameter estimation, we only present results with a demonstrably full-rank FIM in the figures.

## Funding

NIH grants CONNECTS UM1NS132207, S10 RR029672, P41 EB027061 The Geneva Foundation (DoD) S-11229-02

## Author contributions

Conceptualization: H.F., C.L., T.G., Methodology: H.F., Y.C., G.R., T.G., Investigation: H.F., Visualization: H.F., C.L., Supervision: C.L., T.G., Writing—original draft: H.F., Writing—review & editing: C.L., T.G., Y.C., G.R.

## Competing interests

All other authors declare they have no competing interests.

## Data and materials availability

All data needed to evaluate the conclusions in the paper are present in the paper and/or the Supplementary Materials. The codes for the TensorFlow implementation of biophysical models and computing CRLB will be made available on the CMRR website and GitHub.

## Supplementary Information

### Supplementary Text

#### Cramér-Rao lower bound (CRLB) Derivation with Gaussian Noise Assumption

Let us model predicted MR signal attenuation as *ŷ* = *f*_*α*,*g*_(*x*), and let *w* be additive Gaussian noise with zero mean and variance = *σ*^2^. Then, the measured MR signal attenuation *y* is defined as:

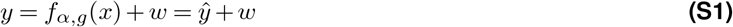

We define the likelihood function function with the Gaussian noise assumption:

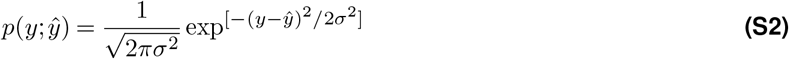

where *E*(*ŷ*) = *y* and 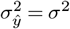

The log-likelihood of Eq. (S2) is

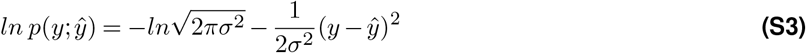

Its first derivative with respect to *x* is

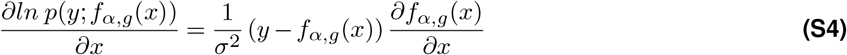

And the second derivative with respect to *x* is

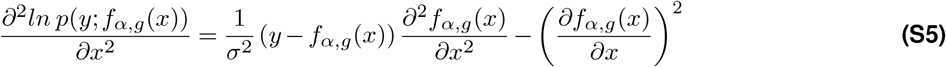

Taking the expected value of Eq. (S5):

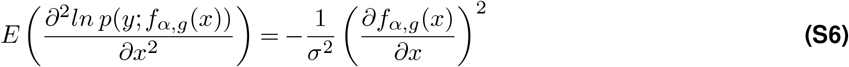

The Fisher information matrix *F*

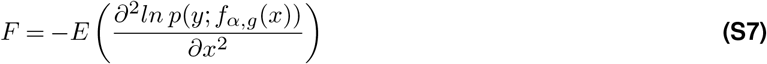

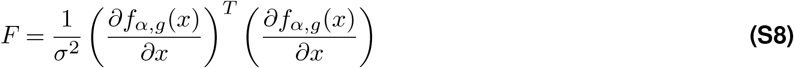

*F* is full rank because of 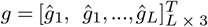 We aim to minimize the sum of the coefficients of variation of the model parameters x. To optimize acquisition parameters *α*, the sum of standard errors of each model parameter *x* is minimized :

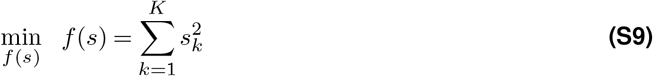

where 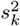 is the variance of *k*th model parameter. Since *s*_*k*_ are unknown, CRLBs are used in their place. CRLBs are estimate by using the inverse of the Fisher information matrix, *F* :

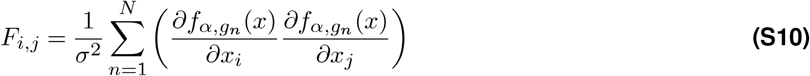

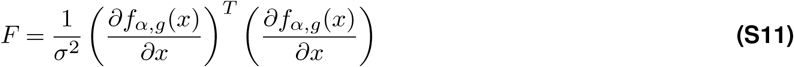

Therefore. the objective function can be written as:

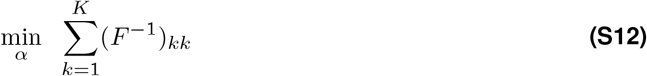

And the CRLB for model parameters (*x*) are diagonal elements of *F*^−1^.

**Figure S1.**
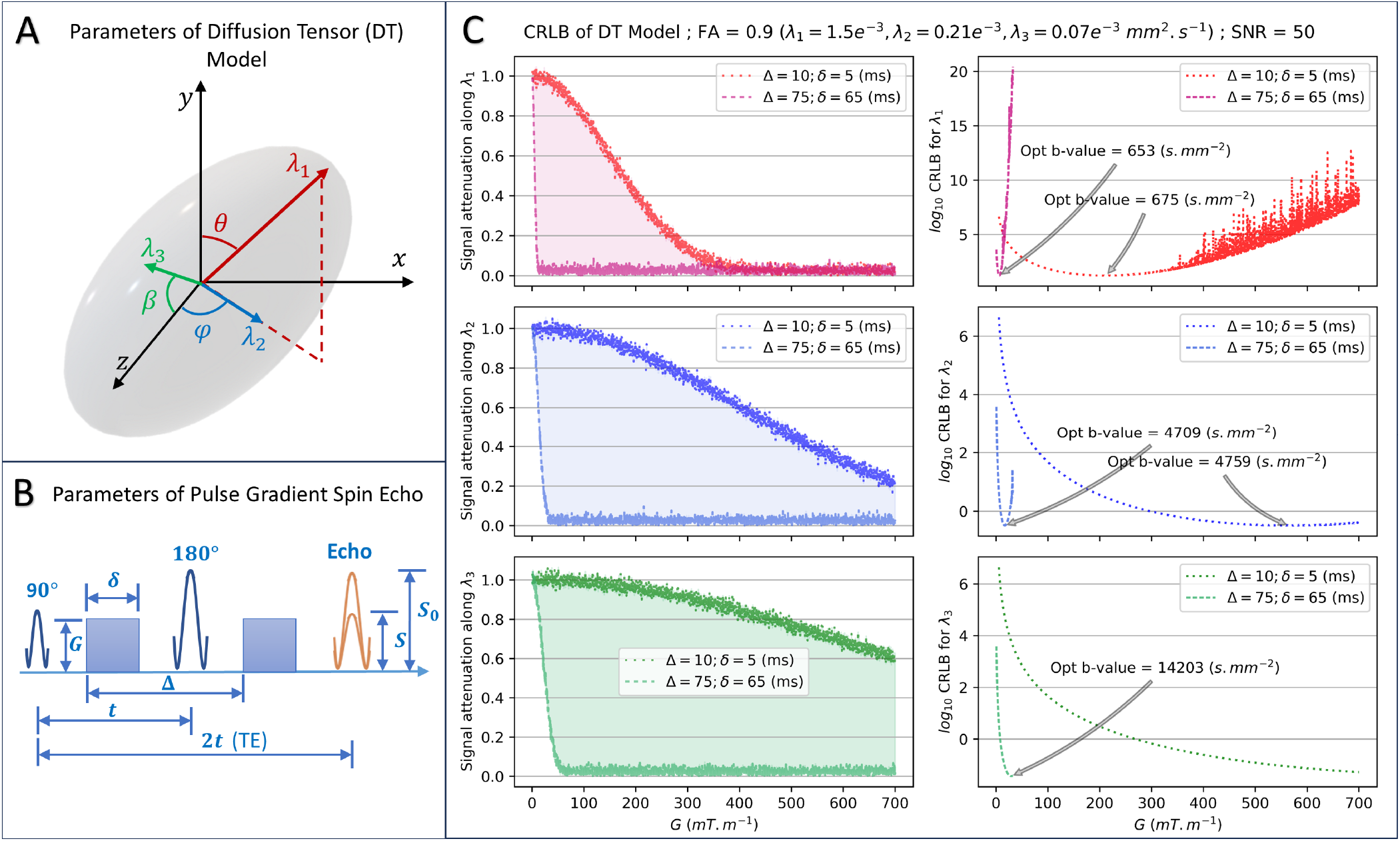
Parameters of the diffusion tensor model and using PGSE sequence. (**A**) Parameters of the diffusion tensor model include the principal eigenvalue *λ*_1_, secondary eigenvalue *λ*_2_, and tertiary eigenvalue *λ*_3_ (*mm*^2^.*s*^−1^), represented in red, blue, and green colors, respectively. The corresponding orientation angles *θ, ϕ*, and *β* (*rad*) are also shown, using the same color code. (**B**) PGSE parameters, specifically the amplitude of the diffusion gradient *G* (*mT*.*m*^−1^), separation time, Δ (*ms*), and duration time, *δ* (*ms*) of the diffusion encoding gradients, can be chosen to achieve optimal experimental design in dMRI-based biophysical modeling. (**C**) The three rows depict outcomes related to the three eigenvalues of the diffusion tensor, following the color code from panel A. The middle column demonstrates the attenuation of the dMRI signal with increasing *G* values, while the right column presents the *log*_10_CRLB plots. Both columns show results for the two extreme combinations of Δ and *δ*. The results show that the minimum CRLB is linked to b-values approximately reciprocal to the corresponding diffusivity values. This implies that higher diffusivity values yield optimal CRLB at lower b-values, while lower diffusivities require higher b-values for reliable estimation.

**Figure S2.**
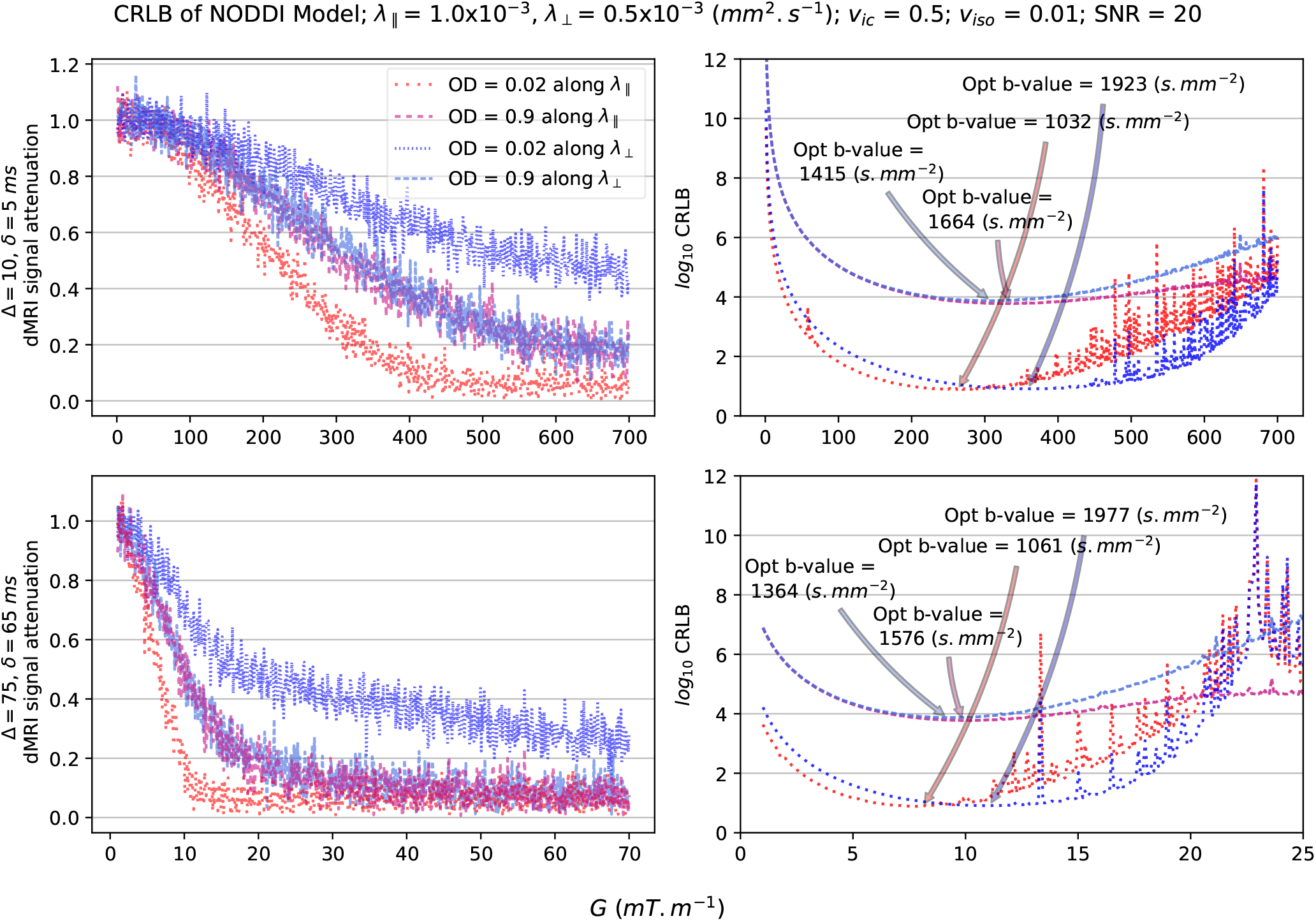
CRLB for lower intrinsic and perpendicular diffusivity values e.g., in gray matter and infant brains using NODDI model. The left column demonstrates the dMRI signal attenuation with increasing *G* values, while the right column presents the log-scale CRLB plots. Dashed/dotted lines show results related to extreme values of *OD* as described in the figure legend. Also, each row shows results for the two extreme combinations of Δ and *δ* (*ms*). The results show that the minimum (optimal) CRLB is related to b-values approximately reciprocal to the corresponding diffusivity values while OD values affect the relationship.

**Figure S3.**
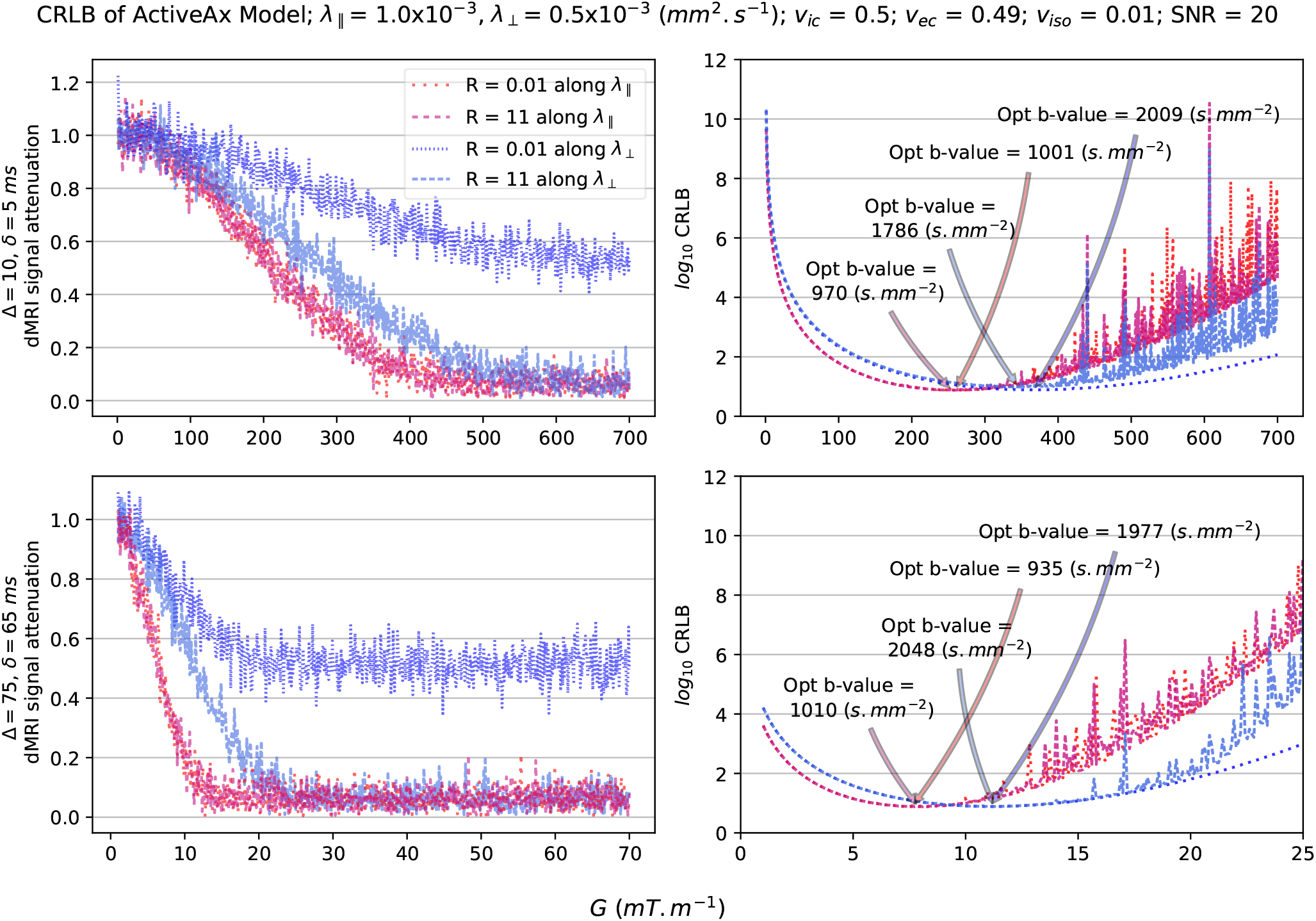
CRLB for lower intrinsic and perpendicular diffusivity values e.g., in gray matter and infant brains using ActiveAx model. The left column demonstrates the dMRI signal attenuation with increasing *G* values, while the right column presents the log-scale CRLB plots. Dashed/dotted lines show results related to extreme values of *R* as described in the figure legend. Also, each row shows results for the two extreme combinations of Δ and *δ* (*ms*). The key finding is that the minimum (optimal) CRLB is related to the b-values approximately reciprocal to the corresponding diffusivity values while *R* values. However, the *R* value only influences this relationship for perpendicular diffusivity *λ*_⊥_.

**Figure S4.**
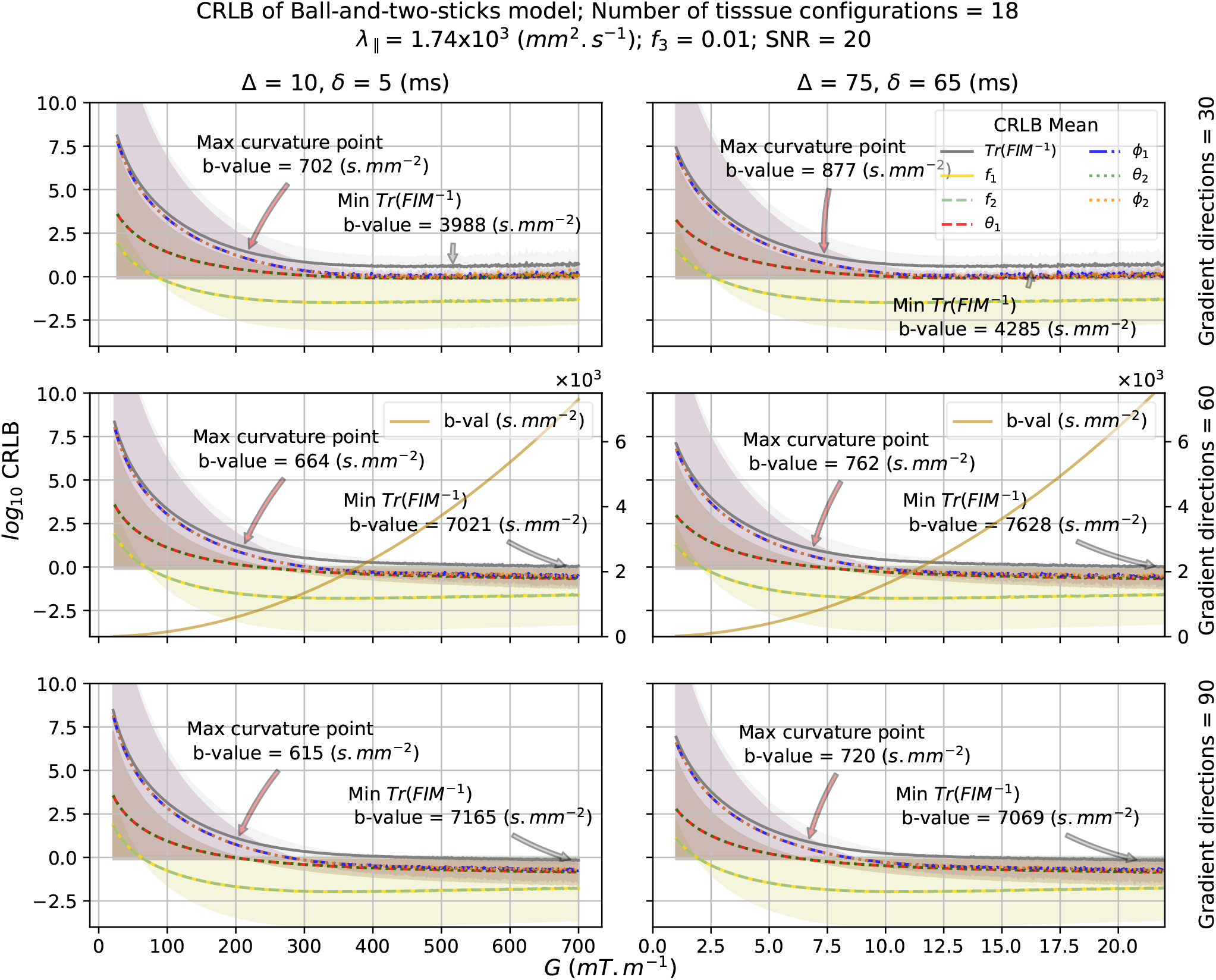
CRLB for Ball-and-Two-Sticks model. CRLB using 18 unique tissue configurations at SNR = 20 are shown. Ball and Stick parameters were constrained for the human brain’s feasible range. The sub-figures share a common y-axis, showing log-scale CRLB plots for all six variables in the model. The x-axis shows the magnitude of the gradient value *G* (*mT*.*m*^−1^) with each columns featuring different G ranges, reflecting variations in the choice of Δ and *δ* (*ms*). b-values corresponding to the optimal CRLB are shown. Sub-figures show plots for mean CRLB values of model parameters and their sum (trace of the CRLB matrix), with the corresponding-colored shaded areas indicating the 95 percent confidence intervals. Maximum curvature points shown with red arrows are “sweet spots” detected by the ‘kneedle’ algorithm with default settings. The points show optimal trade off at which the relative cost of increasing G or b-values is no longer worth for improvement of CRLB.

## Notes

### Competing Interest Statement

The authors have declared no competing interest.

